# SIRT6 is a DNA Double-Strand Break Sensor

**DOI:** 10.1101/765172

**Authors:** Lior Onn, Miguel Portillo, Stefan Ilic, Gal Cleitman, Daniel Stein, Shai Kaluski, Ido Shirat, Zeev Slobodnik, Monica Einav, Fabian Erdel, Barak Akabayov, Debra Toiber

## Abstract

DNA double strand breaks are the most deleterious type of DNA damage. In this work, we show that SIRT6 directly recognizes DNA damage through a tunnel-like structure, with high affinity for double strand breaks. It relocates to sites of damage independently of signalling and known sensors and activates downstream signalling cascades for double strand break repair by triggering ATM recruitment, H2AX phosphorylation and the recruitment of proteins of the Homologous Recombination and Non-Homologous End Joining pathways. Our findings indicate that SIRT6 plays a previously uncharacterized role as DNA damage sensor, which is critical for initiating the DNA damage response (DDR). Moreover, other Sirtuins share some DSB binding capacity and DDR activation. SIRT6 activates the DDR, before the repair pathway is chosen, and prevents genomic instability. Our findings place SIRT6 at the top of the DDR and pave the road to dissect the contributions of distinct double strand break sensors in downstream signalling.

## Introduction

DNA safekeeping is one of the most important functions of the cell, allowing the transfer of unchanged genetic material to the next generation, as well as the proper cellular functioning. Therefore, cells have evolved a sophisticated array of mechanisms to counteract daily endogenous and environmental assaults on the genome. These mechanisms rely on the recognition of the damaged DNA and its subsequent signalling. This signalling cascade triggers responses such as checkpoint activation and energy expenditure and initiates the DNA repair process (Bartek and Lukas, 2007; Bartek et al., 2002; Ciccia and Elledge, 2010; Filippo et al., 2008; Hoeijmakers, 2009; Iyama and Wilson, 2013; Jackson and Bartek, 2009; Lieber, 2008; Madabhushi et al., 2014). If DNA damage is not properly recognized, all the downstream signalling will be impaired. Among the various types of DNA damage, the most deleterious are double-strand breaks (DSBs), which can cause translocations and the loss of genomic material. Until now, very few DSB sensors have been identified, among them Poly ADP-ribose polymerase-1(PARP), MRN complex (MRE11, NBS1, RAD50) and Ku70/80. All of them initiate downstream signalling cascades which usually lead to the activation of specific repair pathways such as Homologous Recombination (HR) or Classical Non-Homologous End Joining (C-NHEJ) (Andres et al., 2015; Sung et al., 2014; Woods et al., 2015). However, how a pathway is chosen is not fully understood, but it is known that the identity of the DSB sensor influences the outcome. For example, MRN complex is associated to HR, while Ku70/80 to NHEJ. Once DNA damage is recognized, transducers from the phosphoinositide 3-kinase family (e.g., ATM, ATR, and DNA-PK) are recruited to the sites of damage. They initiate a broad cascade, recruiting and activating hundreds of proteins which regulate the cellular response including cell cycle progression, transcription, and metabolism. Ultimately, this response will determine whether the cell would live, senesce, or die. Failure to recognize and repair DSBs may lead to tissue ageing and disease (Ciccia and Elledge, 2010; Filippo et al., 2008; Gasser et al., 2016; Ribezzo et al., 2016; Shiloh, 2014).

Sirtuin 6 (SIRT6) is a chromatin-bound protein from a family of NAD^+^-dependent deacylases and ADP-ribosylases. Through these functions, SIRT6 regulates DNA damage repair (DDR), telomere maintenance, and gene expression (Feldman et al., 2013; Jiang et al., 2013; Kugel and Mostoslavsky, 2014). The importance of SIRT6 on DNA maintenance is exemplified in SIRT6-KO mice phenotypes, which include accelerated ageing, cancer and neurodegeneration (Kaluski et al., 2017; Stein and Toiber, 2017; Tasselli et al., 2017; Zorrilla-Zubilete et al., 2018; Zwaans and Lombard, 2014). SIRT6-deficient cells exhibit genomic instability, increased aerobic glycolysis and defects in DNA repair, among other phenotypes (Kugel and Mostoslavsky, 2014; Stein and Toiber, 2017; Tasselli et al., 2017). Moreover, it was recently shown that the capacity of SIRT6 to repair DSB, but not NER, is directly linked to longevity (Tian et al., 2019). We have previously shown that SIRT6 is one of the earliest factors recruited to DSBs, arriving at the damage site within 5 seconds, and allowing the opening of chromatin at these sites by recruiting the chromatin remodeler SNF2H (Toiber et al., 2013). In addition, the silencing of SIRT6 resulted in impaired downstream signalling, affecting the recruitment of key repair proteins such as Ku80, BRCA1 and 53BP1, among others, which are involved in both NHEJ and HR (Daley and Sung, 2014; Bunting et al., 2010; Chen et al., 2017; Escribano-Díaz et al., 2013; Gupta et al., 2014; Kaidi et al., 2010; McCord et al., 2009; Tang et al., 2013; Toiber et al., 2013). These studies indicate that SIRT6 plays important roles at very early stages of the DDR, revealing its key role in DNA repair.

## Results

### SIRT6 arrives to sites of damage independently from other sensors or signaling

The prominent role of SIRT6 in the early steps of DNA damage signalling raises the fascinating possibility that it is also directly involved in DSB sensing. In this work, we set out to study this hypothesis. Firstly, we set to investigate the relationship between SIRT6 and the three known DSB sensors, PARP1, MRE11 (of MRN complex), and Ku80 (of Ku complex).

PARP proteins are among the fastest known enzymes to arrive at DSBs, and their absence is known to impair the recruitment of DSB repair enzymes such as MRE11, NBS1 and Ku80 (Haince et al., 2008; Yang et al., 2018). We inhibited PARP1 by Olaparib and tracked SIRT6 recruitment to sites of laser induced damage by live cell imaging. Interestingly, SIRT6 recruitment was found to be independent of PARP1. SIRT6 arrived at the damage sites even when PARP1 was inhibited, while the recruitment of the macroH2A macro domain, used as a control, depended entirely on PARP1 activity (Fig. 1A-B, Fig S1A-B). Afterwards, we silenced MRE11 and observed impaired NBS1 recruitment, while it had no effect on SIRT6 (Fig. 1C-D, Fig. S1C-D). Ku80 silencing resulted in the expected defective Ku70 recruitment, but did not impair SIRT6 arrival and even higher amounts of SIRT6 were recruited to the site of damage (Fig. 1C-D, Fig. S1E-F). Moreover, when we tested the effect of SIRT6-KO (Fig. S1G) on the recruitment of MRE11 and Ku80, we found that while the former was defective (Fig. 1E-F), Ku80 was not affected by the lack of SIRT6 (Fig. 1G-H). This suggests that SIRT6 is part of MRN DSB recognition, but not of the Ku complex. Next, we silenced ATM and H2AX, both involved in DDR signalling (Fig. S1H). Even though signalling was defective (Fig. S1I-J), SIRT6 arrived to sites of damage independently of these factors (Fig. S1K-L). This indicates that if DSB are present, SIRT6 recruitment is independent of known DSB sensors and DNA damage signalling. Thereafter, we tested whether SIRT6 can be recruited by the initiation of a DNA damage response in the absence of actual DNA damage (lack of DSBs). To answer this question we took advantage of a tethering assay in which we used U2OS cells containing 256x lactose operator (LacO) repeats in their genome (Tang et al., 2013) (Tang et al., 2013) (Tang et al., 2013) and transfected with chimeric proteins containing lactose repressor (LacR) conjugated to known DDR initiating repair enzymes (Scheme in Fig. 2A) (Soutoglou and Misteli, 2008). In this system, ATM (ATM-LacR-Cherry) initiates the DNA damage response by its mere presence on chromatin, shown by H2AX ser-139 phosphorylation (ϫH2AX) (Fig. S2A-B). However, in this system with no actual DNA damage, ATM failed to recruit SIRT6 to LacO site, even though signalling was taking place and H2AX was phosphorylated (Fig. 2B-C). As a control, known interactors such as SNF2H and Ku80 (McCord et al., 2009; Toiber et al., 2013) did recruit SIRT6 to the tethering sites (Fig. 2B-C, Fig. S2C-D). Moreover, MRE11 and NBS1 also recruited SIRT6 to the LacO site (Fig. S2C-D) suggesting that there is either direct interaction between them or they work together in DDR, also indicated by the impaired recruitment of MRE11 to LID.

**Figure 1.**
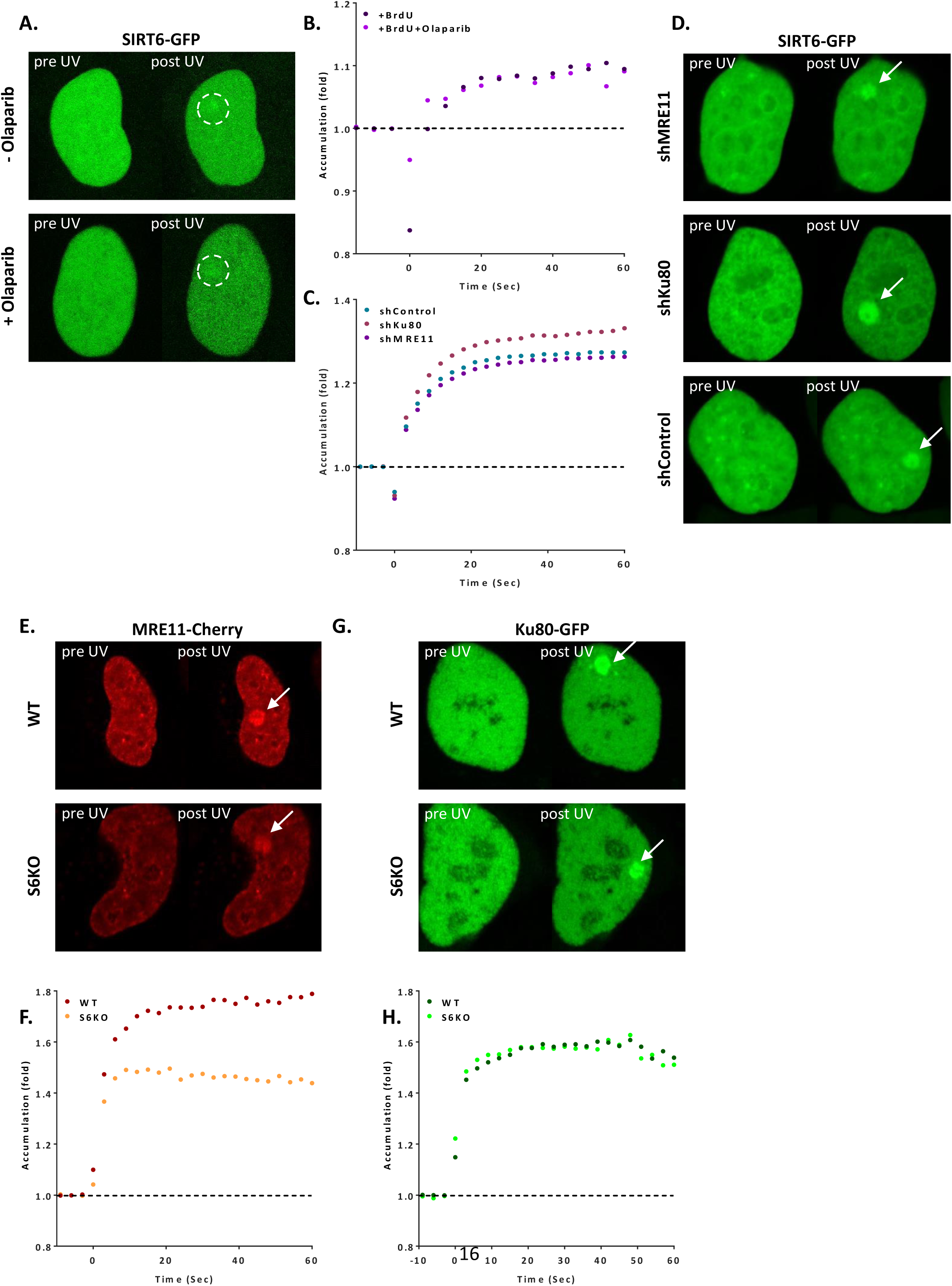
SIRT6 arrives to sites of damage independently of other repair enzymes. **(A-B)** Live imaging for SIRT6-GFP upon laserinduced damage (LID) in naïve U2OS cells treated with BrdU and Olaparib (n=10) and cells treated with BrdU alone (n=10), 5sec intervals. **(C-D)** Accumulation of SIRT6-GFP at sites of LID in control cells (n=38) or shMRE11 (n=40) or shKU80 (n=38), at 3sec intervals. **(E-H)** Accumulation of MRE11-Cherry in SIRT6KO (n=16) or WT cells (n=20). 3sec intervals. Or accumulation of KU80-GFP at LID sites in SIRT6KO (n=17) or WT (n=17) cells at 3sec interval.

**Figure 2.**
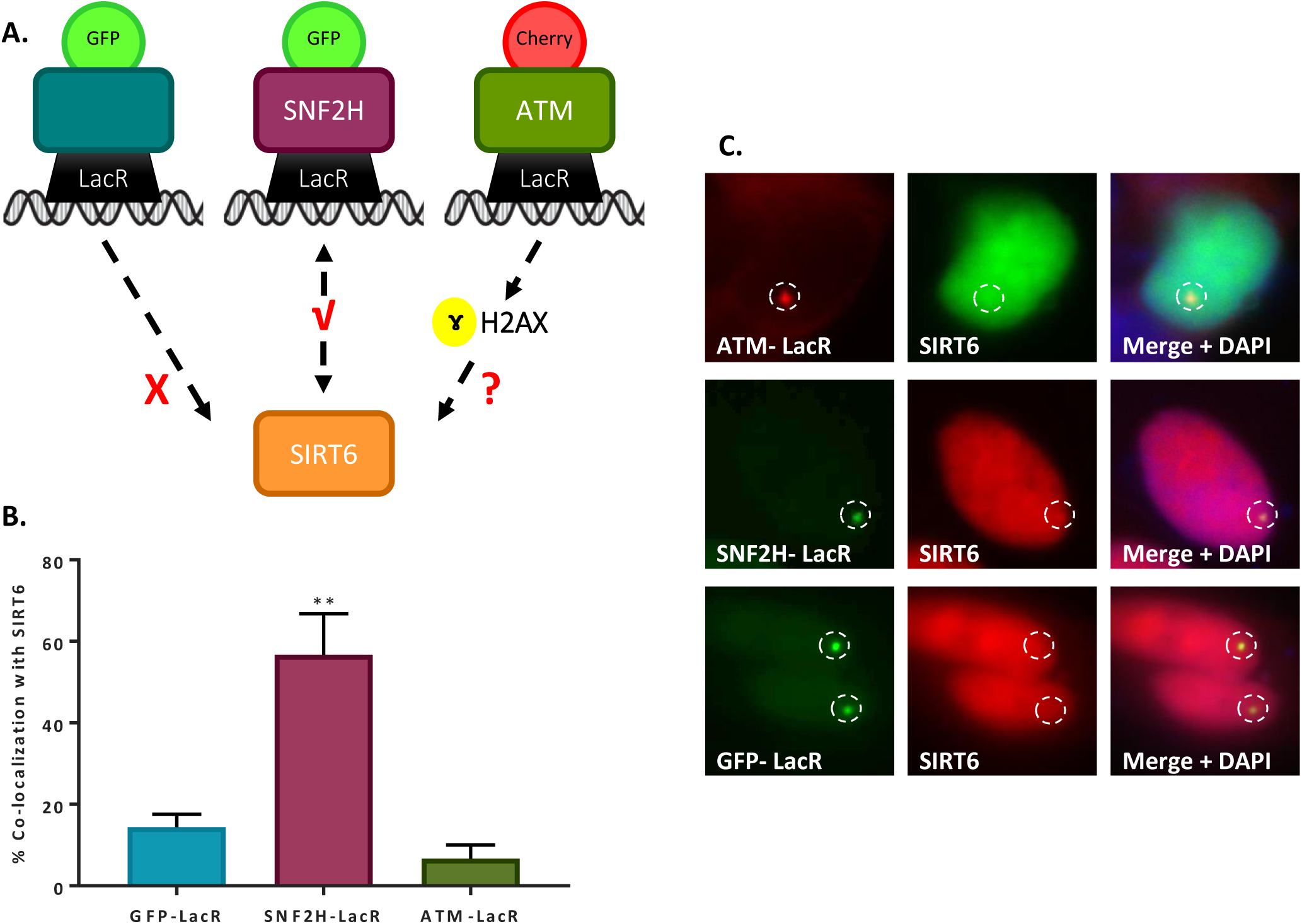
SIRT6 is not recruited by signaling alone. **(A)** Schematic representation of the “Tethering assay” Recruitment can occur through DDR signaling (LacR-ATM-Cherry) or through direct protein-protein interaction (LacR-SNF2H-GFP). **(B-C)** Recruitment of SIRT6-GFP/SIRT6-Cherry to LacO sites by LacR-ATM Cherry (n=30, p>0.05), LacR-SNF2H GFP (n=85, p<0.005) and LacR-GFP (n=85). The graph depicts averages of 3-6 experiments +/- SEM.

Taken together, these results indicate that SIRT6 arrives to the sites of damage independently of other known repair sensors and enzymes, and that in the absence of an actual DSB or damaged DNA, signalling itself is not sufficient to bring SIRT6 to the damage sites.

### SIRT6 directly binds DNA DSBs

The findings described so far suggest that SIRT6 responds selectively to the actual damage, and that silencing or inhibiting key factors in DDR do not affect its recruitment. Therefore, we tested whether SIRT6 could detect the actual DNA break itself. We first measured SIRT6 capacity to bind naked DNA by Electrophoretic Mobility Shift Assay (EMSA). We found that SIRT6 was able to bind naked DNA without preference for a sequence (we tested different oligos and restricted sites) (Fig. 3A-B, S3A). We studied SIRT6’s preference for several DNA damage structures, including dsDNA with blunt or overhanging ends as well as RNA. SIRT6 has the ability to bind them all, still, it binds RNA with much lower affinity (Fig. S3B). SIRT6 exhibits the highest affinity towards ssDNA (Kd=1.39μM) which is similar to the values for MRE11 (Kd∼1μM) (Williams et al., 2008) and Ku80 (Kd=0.4μM) (Arosio et al., 2002). Interestingly, based on the curve fitting, SIRT6 seems to bind ssDNA at one site as a monomer. In contrast, there seem to be multiple binding sites on a blunt and sticky-end DNA (Hill Slope greater than 1), suggesting that for open ended dsDNA each molecule of SIRT6 can bind one DNA strand (Fig. 3A-B, S3A, scheme in Fig. 3C). Since in EMSA all the DNA used had open ends, we developed an additional DNA binding assay based on the co-immuno-precipitation of a plasmid (IP-qPCR). In brief, flag-tagged repair proteins were purified and incubated with DNA, then immuno-precipitated along with the DNA that they bound. The DNA was later purified and its enrichment was measured by qPCR. Proteins were incubated either with a circular plasmid or the same plasmid presenting blunt or sticky ends. As expected, NBS1, which does not bind DNA by itself (Myler et al., 2017), did not bind either plasmid (open or closed ends). In contrast, SIRT6 and MRE11 had high affinity to liner DNA, however, they showed almost no binding for closed plasmids (Fig. S3C). Moreover, SIRT6 exhibited a higher affinity to sticky ends over blunt ends (Fig. 3D), a structure that has a high resemblance to DSBs. In addition, it did not distinguish between 3’ or 5’ overhangs (Fig. S3D). These assays indicate that SIRT6 does not function by binding intact DNA or a particular sequence, but rather by binding to open DNA ends, and particularly, to ssDNA. It is important to note that this capacity is independent of the presence of NAD^+^, the known cofactor of SIRT6 (Fig. 3E), and does not activate SIRT6 catalytic activity (Fig S3E). Moreover, SIRT6 was able to protect the open ends of DNA from exonuclease activity (ExoI) preventing the access to exonuclease cleavage, just as in the case of MRE11, implying that SIRT6 specifically binds to DNA ends (Fig. S3F-G).

**Figure 3.**
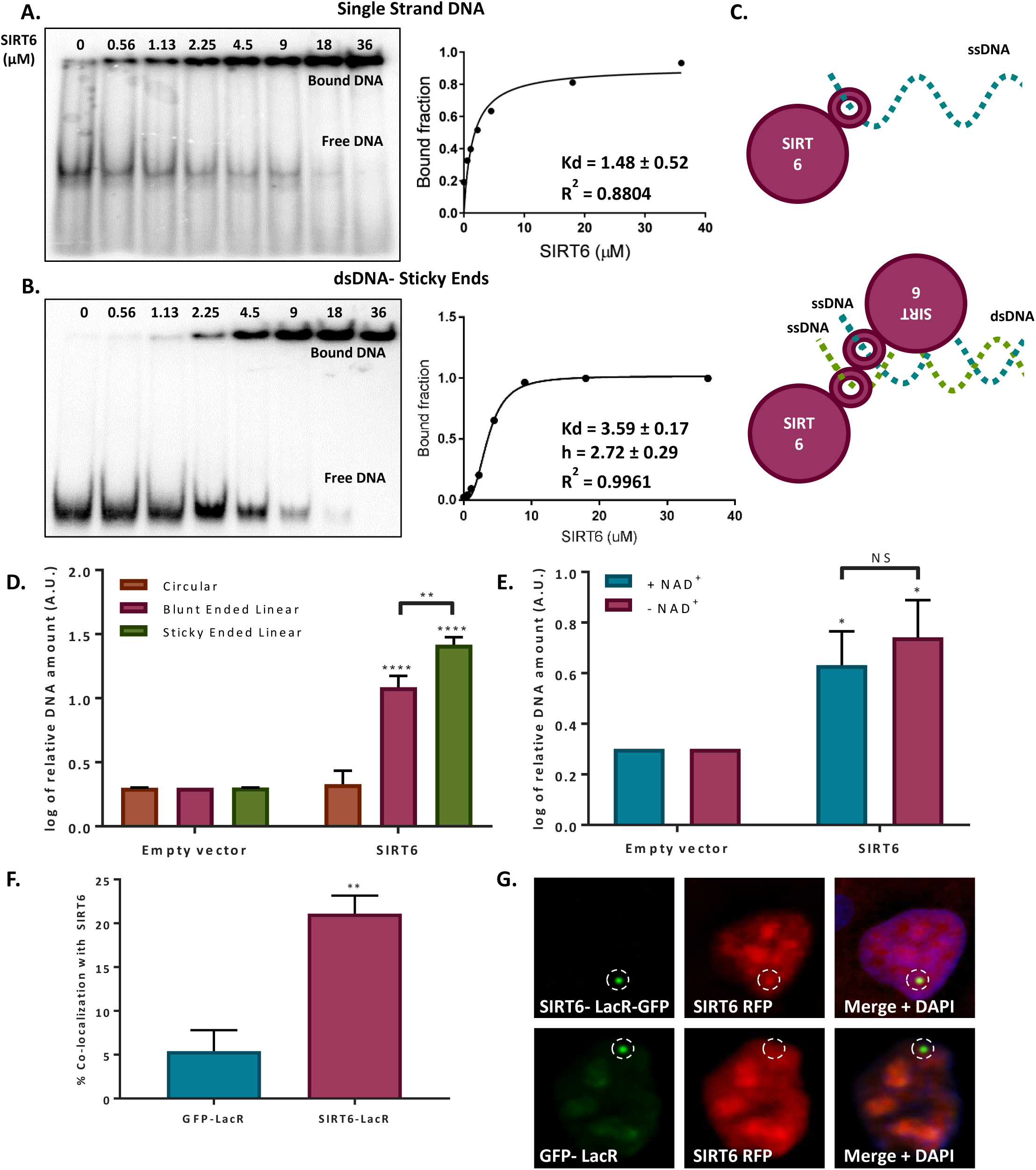
SIRT6 binds DNA with no intermediates. **(A-B)** Gel retardation assay of 32P-5’ end-labeled single strand DNA and sticky ended dsDNA as a function of increasing concentrations of SIRT6-His. Kd value (ssDNA: Kd= 1.39 ± 0.53, sticky dsDNA: Kd=4.52 ± 0.09). **(C)** Suggested model of SIRT6 binding ssDNA as a monomer or open ssDNA ends of dsDNA as a dimer. **(D)** SIRT6-Flag DNA binding ability of circular, blunt ended and sticky ended cleaved plasmids. The graph depicts averages 3 experiments +/- SEM, after logarithmic transformation. **(E)** SIRT6-Flag DNA binding ability of an open-ended plasmid with and without NAD^+^. Averages of 4 experiments +/- SEM, after logarithmic transformation. **(F-G)** Dimerization of SIRT6 at LacO site, represented by the recruitment of SIRT6-Cherry by LacR-SIRT6-GFP (n=181, p<0.005) or LacR-GFP (n=104). The graph depicts averages of 4 experiments +/- SEM.

### SIRT6 binds DNA ends as a dimer

Our EMSA results for WT SIRT6 indicate that SIRT6 binds ssDNA with no cooperativity, suggesting a single binding site. In contrast, when the substrate were dsDNA oligos, we found the Hill coefficient to be greater than one, indicating cooperativity (Fig. 3A-B, S3A). These results suggest that a single molecule of SIRT6 binds ssDNA. Even so, given two ssDNAs, such as open ended DSB, one SIRT6 molecule will interact with another allowing a dimer of SIRT6 to bind a single molecule of dsDNA with two open ends in a single side, 5’ or 3’ (see schematic Fig. 3C). Together, the two SIRT6 molecules show cooperativity. Interestingly, SIRT6 known Crystal Structure presents a dimer conformation (Jiang et al., 2013; You et al., 2017). To further characterize the structure of SIRT6 in a solution, we used Size Exclusion Chromatography-Multi-Angle Light Scattering (SEC-MALS) and Small-Angle X-ray Scattering (SAXS). Importantly, both methods showed that SIRT6 tends to aggregate, however, by using SEC-MALS we noted that the aggregation was significantly reduced by the presence of DNA oligomers (Fig. S3H), which suggests that SIRT6 is stabilized by and favours DNA interactions. SAXS data provides a low-resolution structure of SIRT6, presumably corresponding to a tetramer (Fig. S3I-K), supporting the model suggested by the EMSA results (with dimers at the 5’ and 3’, a tetramer). The result obtained by SAXS does not exclude the presence of SIRT6 dimers or trimers in solution. Lastly, we measured dimerization in vivo by taking advantage of SIRT6-LacR-GFP localization at LacO sites, as well as the recruitment of SIRT6-RFP, observing a significant co-localization percentage of both SIRT6 molecules (Fig 3F-G).

Overall, our predictions suggest that SIRT6-DNA complex is organized in dimers, probably at each end of the DNA oligomers. Moreover, based on the reconstructed SAXS structure we can observe a compaction of SIRT6 in the presence of DNA suggesting a conformational change (Fig. S3I-K).

### SIRT6 binds ssDNA through its core domain, that forms a “tunnel-like” structure

As SIRT6 has not been previously reported in the literature to be a DNA binding protein, we aimed to identify the domain involved in ssDNA binding. To this end, we first analyzed SIRT6 structure to find a potential DNA binding domain. We found a region within the core domain (28 a.a.) that had potential for binding DNA (Fig. 4A-C). We purified full length SIRT6 (SIRT6 FL) and a fragment of the core domain alone (Core: from a.a. 34 to 274). Both of them were able to bind DNA with similar affinities, indicating that the core domain is the main responsible for DNA binding (Fig. 4D). To understand which amino acids could be involved in the DSB binding we mapped them to the known structure of SIRT6 (http://dnabind.szialab.org/). The model points to a subset of amino acids that have a higher likelihood to be involved in DNA binding. Surprisingly, these amino acids are concentrated near a physical structure which resembles a tunnel (Fig. 4A). This tunnel, which has not been previously discussed in the literature, is narrow and could accommodate ssDNA (Fig. 4E), but not larger dsDNA. Lacking an open end, normal undamaged DNA could not enter this tunnel, however, broken DNA ends could. Therefore, we hypothesized that the destruction or disruption of the tunnel will impair SIRT6 DNA binding capacity. To test this hypothesis, we generated several point mutations of the amino acids in the tunnel-like structure of SIRT6 (Fig. S4A-B). Purified SIRT6-MBP point-mutants were tested by EMSA to estimate their DNA binding ability. As predicted, single point mutations in key amino acids at the tunnel impaired SIRT6 binding capacity (including the catalytic dead mutant H133Y) (Fig. 4F-G). The only mutant that showed no effect on binding was D63Y, where the amino acid did not impair the charge as strongly as the D63H. Interestingly, mutations in D63 were previously reported to provoke the loss of SIRT6 function in cancer, and have been recently shown to be lethal in humans (Ferrer et al., 2018).

**Figure 4.**
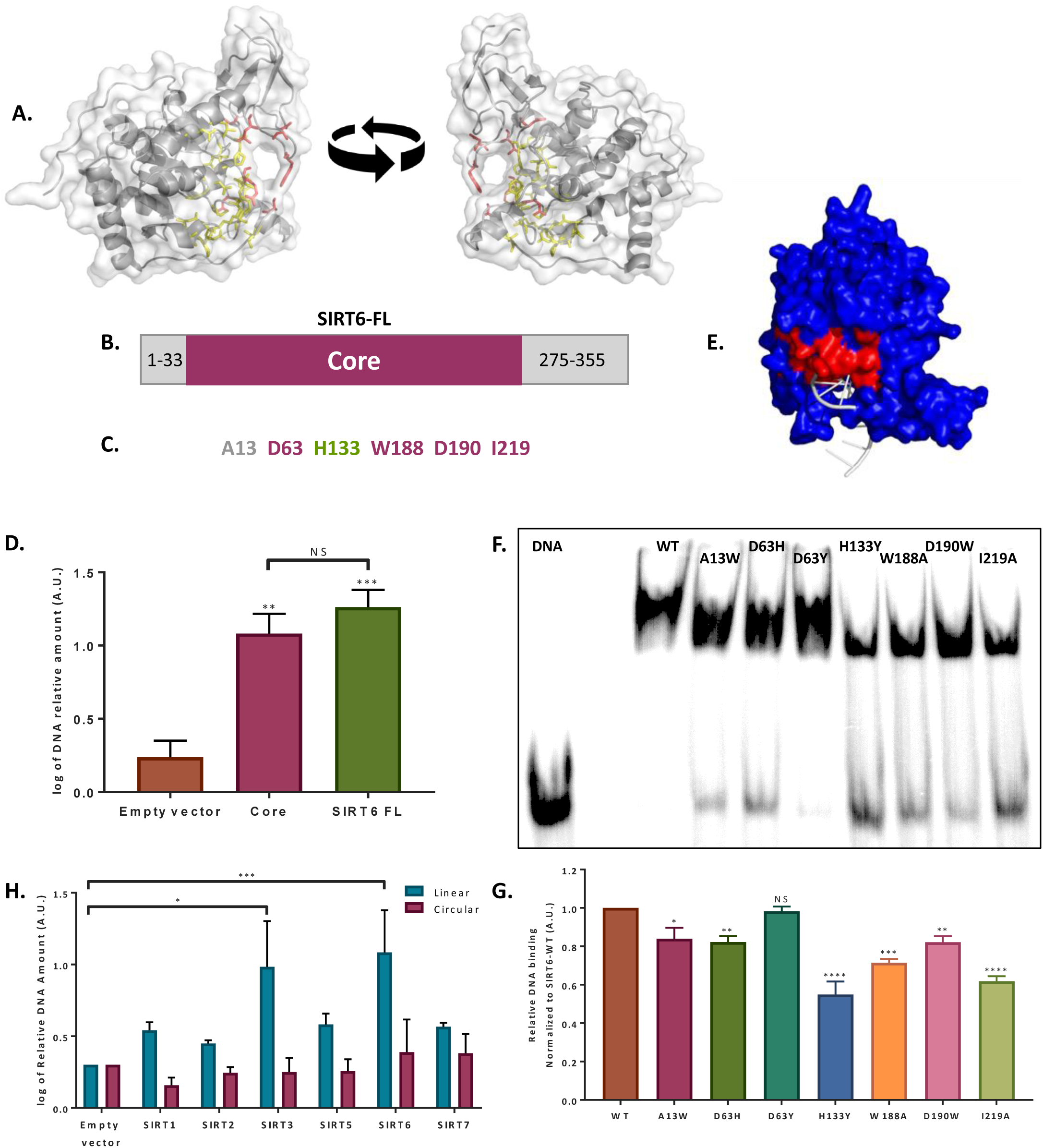
SIRT6 binds DSB through its Core domain. **(A)** DNA binding predicted site based on the published SIRT6 structure, (http://dnabind.szialab.org/). Highlighted in yellow are predicted DNA binding amino acids in the SIRT6 core domain, in red the tunnel forming amino acids that were mutated. **(B)** Schematic representation of SIRT6 core domain. **(C)** List of amino acids that are predicted to participate in the “tunnel-like” structure. **(D)** DNA binding of an open-ended plasmid by full length SIRT6 (p<0.0005) and SIRT6-core domain (p<0.005, log of averages of 3 experiments +/- SEM). **(E)** SIRT6 ssDNA binding prediction, based on the known SIRT6 structure with ssDNA. **(F-G)** Gel retardation assay of 32P-5’ end-labeled single strand DNA with SIRT6-MBP mutants. The graph depicts average +/- SEM of 3 experiments. **(H)** Flag tagged mammalian Sirtuins DNA binding ability of circular and linear plasmids. The graph depicts log of averages of 4-7 experiments +/- SEM. (*=p<0.05, **=p<0.005, ***=p<0.0005)

Since we hypothesized that SIRT6 DNA binding domain is in close proximity to its catalytic domain, we set to examine how these mutations would affect SIRT6 catalytic activity. We performed a Flor-de-lys assay to assess the mutants’ activity using a H3K9-myristolatted peptide. Most mutants showed a decrease in SIRT6 activity compared to SIRT6-WT, however, A13W mutation showed increased SIRT6 activity (Fig. S4C).

These results suggest that the close proximity of the two domains, the catalytic one and the DNA-binding tunnel, may be linked due to the similarity of the ssDNA and NAD^+^ molecules (ssDNA is a polymer of nucleotides, NAD^+^ consists of two nucleotides joined through their phosphate groups). However, one of the mutants showed a slight decrease in binding (A13W) and an increase in catalytic activity, suggesting that these roles could be partially separated.

### DNA binding ability is conserved among other Sirtuins

The core domain of SIRT6, where its DNA binding domain is located, is conserved among all Sirtuins. Therefore, we tested whether other mammalian Sirtuins could bind DSB as well. Our results indicate that all Sirtuins have some capacity to bind broken ended DNA, but some do it with a significantly lower affinity (Fig. 4H, S4D). Only SIRT7 showed binding capacity towards circular DNA, as previously described (Gil et al., 2013). It is also important to note that we tested mouse (SIRT-Flag) and human SIRT6 (SIRT6-His) and they both bind linear, but not circular, DNA (Fig. 4H, S4D).

### SIRT6 can initiate DNA damage response

As shown above, SIRT6 directly recognizes DNA breaks and arrives to the sites of damage independently of DDR signaling. Nonetheless, DNA damage recognition *per se* cannot activate the DNA damage response. Therefore, we set to examine whether SIRT6 has also the capacity to initiate the DDR through downstream signaling. To that aim, we took advantage of the previously described tethering assay with SIRT6-LacR-GFP/Cherry chimeras. Remarkably, SIRT6 has the same ability to induce the activation of the DNA damage response as MRE11, measured by its capacity to activate the phosphorylation of H2AX at the LacO sites compared to LacR-GFP/Cherry. Interestingly, SIRT6 catalytic mutant, SIRT6-HY, was able to initiate the DNA damage response as well, raising the possibility that SIRT6 DDR initiation capacity is independent of its catalytic activity (Fig. 5A-B).

**Figure 5.**
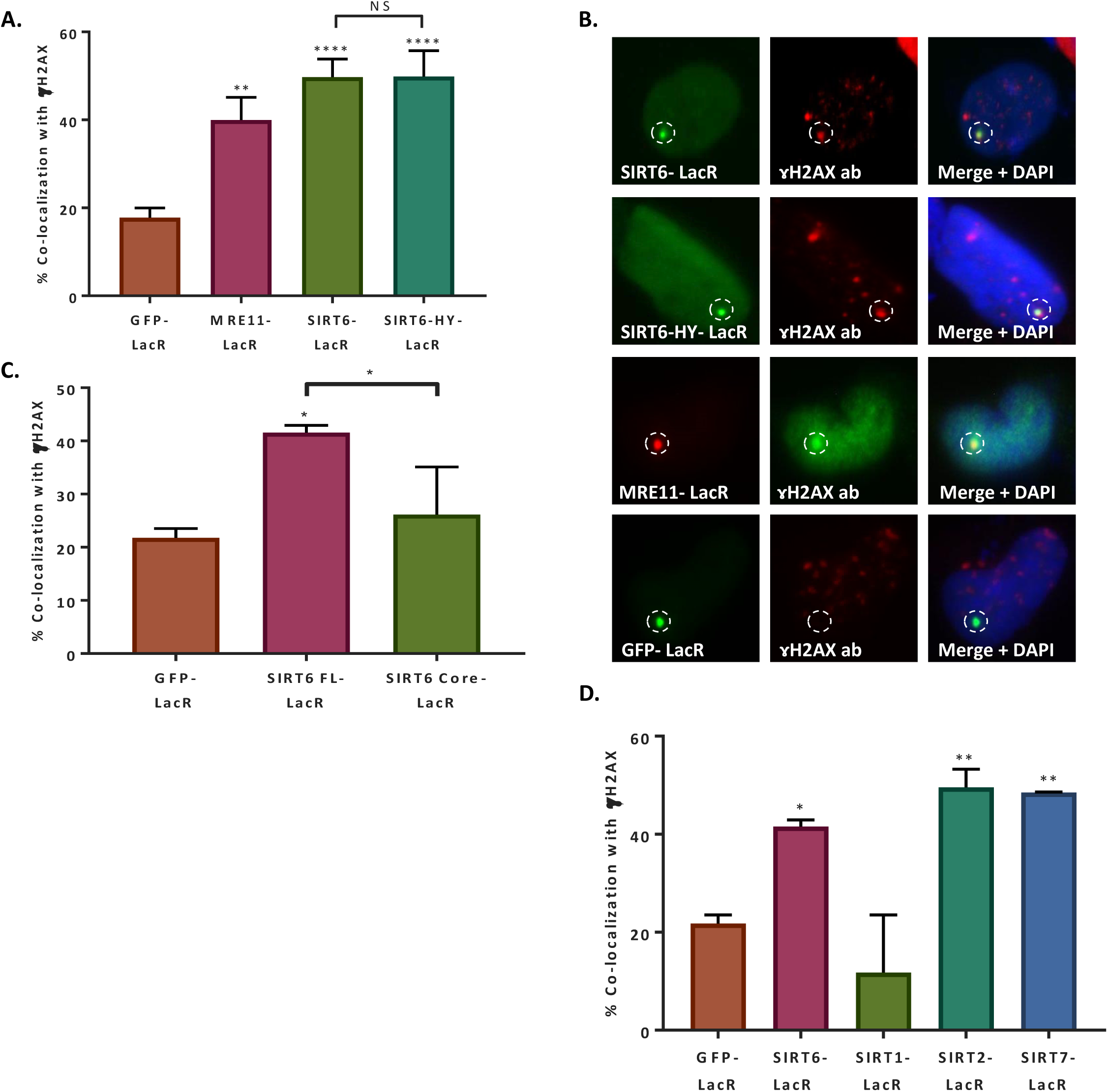
SIRT6 can initiate DNA damage response. **(A-B)** Initiation of DDR was measured by co-localization with ɤh2ax. LacR-GFP (n=310). LacR-MRE11-Cherry (n=136, p<0.005), Lac-SIRT6-GFP (n=243, p<0.0001) or LacR-HY-GFP (n=71, p<0.0001). Averages of 4-9 experiments +/- SEM. **(C)** Initiation of DDR by full length LacR-SIRT6-GFP (n=127, p<0.005), LacR-SIRT6-core –GFP (n=66, p>0.05) or LacR-GFP (n=64). Averages of 3-4 experiments +/- SEM. **(D)** Initiation of DDR LacR-SIRT6-GFP (n=127, p<0. 05), LacR-SIRT1-GFP (n=44, p>0.05), LacR-SIRT2-GFP (n=67, p<0.005), LacR-SIRT7-GFP (n=68, p<0.005) or LacR-GFP (n=64). Averages of 3-10 experiments +/- SEM.

Nevertheless, since SIRT6 can generate dimers, endogenous SIRT6 could dimerizate in the cells with SIRT6-HY-LacR allowing the activation of the DDR. To test this possibility, we used Nicotinamide (NAM) inhibiting this way endogenous SIRT6 (Fig. S5A). However, even when the endogenous SIRT6 was inhibited (shown by the increase in H3K56ac, Fig. S5A), LacR-SIRT6-HY was still able to activate the DDR, supporting the independence of DDR initiation from SIRT6 catalytic activity (Fig. S5B).

It is important to highlight that SIRT6-HY has 50% lower DNA binding capacity to DSB. In fact, it would have failed to bind the DNA and activate the DDR if it was not tethered to chromatin through the LacR domain, bypassing the need of DNA binding and thus allowing SIRT6-HY to initiate the DDR.

To study whether SIRT6 activity and initiation capacity are separate, we tested the Core-LacR-GFP, which has an active catalytic domain but lacks the C and N terminus of SIRT6 (Tennen et al., 2010). We observed that Core-LacR-GFP failed to activate the DDR (Fig. 5C, S5C), suggesting that protein-protein interactions are responsible for DDR signaling through its other domains rather than SIRT6 core domain.

Moreover, we tested LacR-SIRT1, SIRT2 and SIRT7 initiation capacity in the tethering assay since all these Sirtuins have the ability to localize to the nucleus and have been associated with DNA repair (Jeong et al., 2007; Li et al., 2016; Paredes and Chua, 2016; Rifaï et al., 2018; Vazquez et al., 2016, 2017; Zhang et al., 2016). Remarkably, SIRT2 and SIRT7 could initiate the DDR, but SIRT1 could not (see note in methods) (Fig. 5D, S5D). Although other Sirtuins have some binding activity and some initiation capacity, SIRT6 is unique since it has both capacities.

Taken together, these experiments indicate that although SIRT6 binds DNA through its core domain, the activation of downstream signaling does not require the catalytic activity of SIRT6, but its N and C terminus are required for DDR activation.

Lastly, we tested whether SIRT6 could recruit repair factors of the DDR cascade and whether it shows a preference for a certain repair pathway. Although we observed a more prominent effect of SIRT6 on the recruitment of the HR initiator MRE11 rather than the NHEJ initiator Ku80 (Fig. 1E-H), it was previously reported that SIRT6 affects both repair pathways (Chen et al., 2017; Mao et al., 2011; McCord et al., 2009; Tian et al., 2019; Toiber et al., 2013). Indeed, we noticed that SIRT6 deficiency results in impaired recruitment of both 53BP1 and BRCA1 to the sites of laser induced DSBs, suggesting impaired activation of both NHEJ and HR (Fig. S6A).

In order to test SIRT6 ability to recruit these and other DDR factors to the sites of damage, we took advantage of the tethering system once more. Our results show that SIRT6 can recruit proteins from HR such as MRE11, NBS1, ATM and BRCA1, as well as proteins from NHEJ such as Ku80, Ku70 and 53BP1 (Fig. 6A-B). As a control, we tested co-localization with CDT1, a nuclear protein that does not participate in the DDR. As expected, CDT1 was neither recruited by SIRT6 nor by GFP alone.

**Figure 6.**
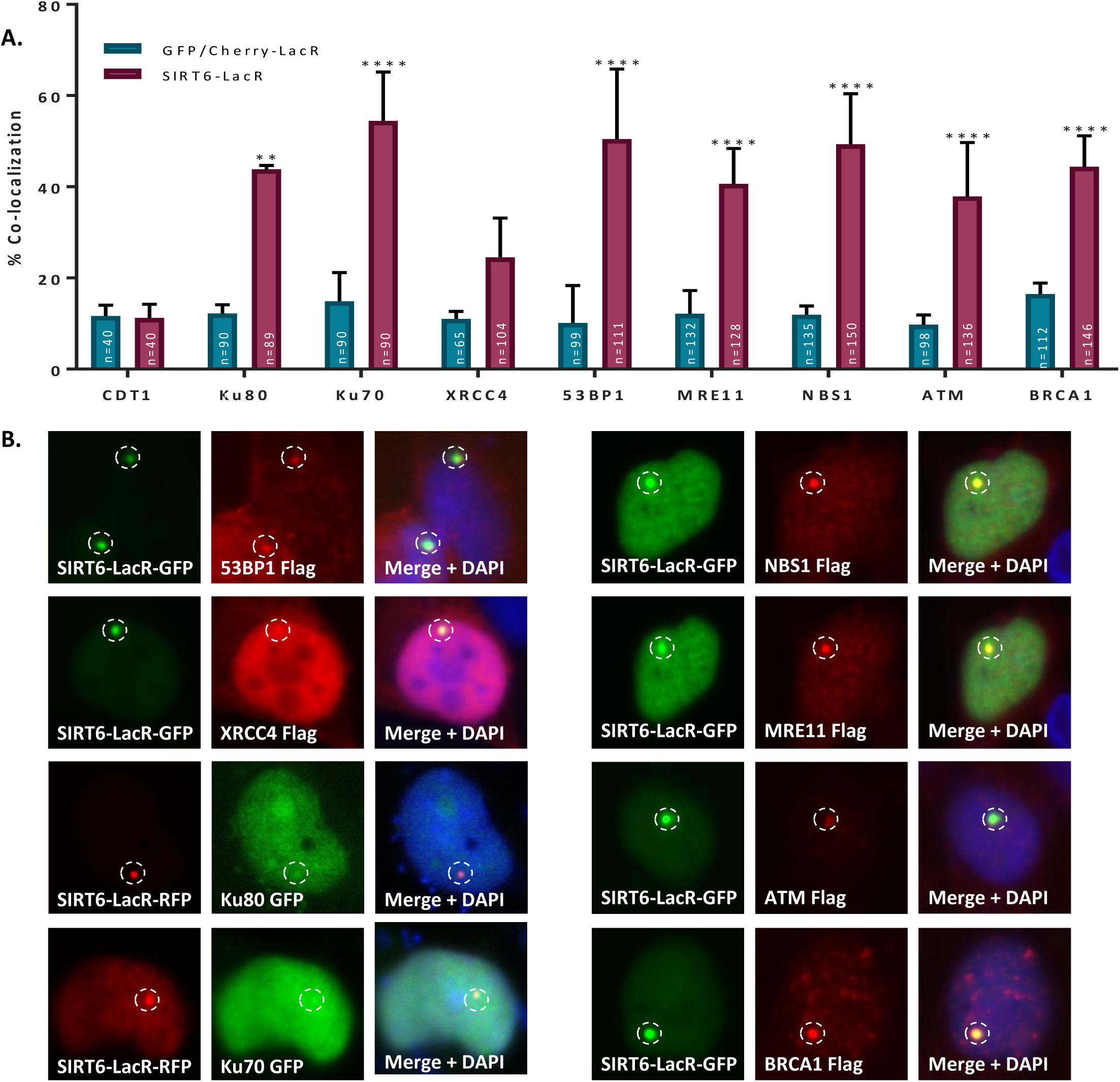
SIRT6 can recruit enzymes of both NHEJ and HR repair pathways. **(A-B)** Percentage of co-localization of repair enzymes with LacR-SIRT6 GFP/LacR-SIRT6 Cherry at LacO sites. IF was done with Flag antibody. The graph depicts averages of 3-6 experiments +/- SEM. (*=p<0.05, **=p<0.005, ***=p<0.0005, ****=p<0.00005).

Since SIRT6 DDR activation is independent of its catalytic activity, we further examined if it is needed for DDR protein recruitment. Taking advantage of the tethering assay, we observed that both SIRT6-WT and SIRT6-HY recruited 53BP1 and BRCA1, meaning that the recruitment is independent of SIRT6 catalytic activity (Fig S6B-C).

53BP1 and BRCA1 can antagonize each other and a change in their concentration within the cell may influence the recruitment capacity. Therefore, we tested through IF whether their overexpression is different from endogenous proteins. However, the results were very similar, suggesting that the recruitment is independent of the amount of protein in the cell, and an additional layer of regulation would influence the recruitment (Fig. S6D-E).

The tethering assay can detect both protein-protein interaction or recruitment through signaling. To differentiate these two possibilities, we inhibited DDR signaling by supplementing the media of the cells with Wortmannin, thus inhibiting ATM, ATR and DNA-PKc (Scheme in Fig. S6F). Our results indicate that when these kinases are inhibited (showed by a reduction in ɤH2AX levels), the recruitment of both 53BP1 and BRCA1 to the LacO site is reduced (Fig. S6G-I). However, the recruitment of the DDR initiators-Ku80 and MRE11 is not affected by Wortmannin, suggesting that their recruitment is based on protein-protein interactions and not on signaling alone (Fig. S6J). These results indicated that DDR initiation by SIRT6 starts with complex formation, leading to the recruitment of transducer proteins and resulting in protein recruitment through signaling.

## Discussion

In this work, we have discovered a novel function for the chromatin factor SIRT6 as a DSB sensor, by which it is able to bind DSBs and initiate the cellular DDR.

We showed that SIRT6 can bind DNA with high affinity for ssDNA and open-ended dsDNA. We believe that the binding occurs through a previously unrecognized tunnel-like structure in the protein core domain, close to its catalytic site. This structure could only fit ssDNA, and while other proteins require resection for ssDNA identification, 3-4 bases are enough for SIRT6.

By generating several point mutations in the hypothesized DNA binding site, we managed to reduce SIRT6 DNA binding capacity, also reducing the catalytic activity of proteins. However, A13W and D63Y mutations raise the possibility that, despite the proximity of these sites, these abilities are distinct ones. While D63Y mutation had no effect on DNA binding, it causes a significant reduction in SIRT6 catalytic activity. A13W mutation, on the other hand, resulted in an increase in catalytic activity along with a slight reduction in DNA binding.

In addition, we showed that SIRT6 can arrive at the sites of DSBs independently of the known sensors and activate the DDR by itself. We also observed that its catalytic activity is not necessary for DDR initiation when it is already bound to the DNA, shown by the ability of SIRT6-HY-LacR to initiate the DDR as well. However, since the binding capacity in the HY mutant is reduced, we believe that SIRT6-HY would not be able to bind and remain in the DNA, and therefore, all DDR initiation would be impaired. Interestingly, even though the initiation of the DDR occurs when SIRT6 is catalytically inactive, it cannot be initiated by the active core-domain alone. These results suggest a complex relationship between binding capacity and activation, in which binding per se cannot result in DDR signaling.

Given that the core domain, which contains both the catalytic and the DNA binding domains of SIRT6, is conserved among Sirtuins, we have also shown that other Sirtuins share the ssDNA binding capacity but with different affinities. This is especially interesting since Sirtuins are present in the cell at different cellular locations (cytoplasm, nucleus and mitochondria) and have different catalytic activities (Deacetylases, deacylases, and ADP ribosylases) (Liszt et al., 2005). This suggests that the DSB binding capacity could be relevant in other cellular compartments, such as in mitochondrial DNA repair. When nuclear SIRT2 and SIRT7 were forced to localize to the DNA by the LacO-LacR tethering assay, they were able to initiate the DDR as well. However, SIRT7 lacks the broken DNA binding specificity and SIRT2 has a poor binding capacity, which indicates that their sensing capacity is less critical for the cell as exemplified as well in the KO mouse models (Kim et al., 2011; Shin et al., 2013).

These findings open new possibilities for the cellular functions of the Sirtuin family; nevertheless, we believe in the uniqueness of SIRT6 since it possesses all these abilities at once.

Putting SIRT6 at the top of DSB repair might explain why the lack of SIRT6 gives place to one of the most striking phenotypes in human, monkeys and mice, including typical phenotypes with genomic instability such as premature ageing, accelerated neurodegeneration, tissue atrophy and cancer (Kugel and Mostoslavsky, 2014; Tasselli et al., 2017). In particular, SIRT6 is involved in several repair pathways. As a sensor and DDR initiator, its absence would have deleterious effects on the whole downstream DDR signaling. Our results point out that its role begins as a DSB sensor (although it may recognize other DNA lesions), recognizing and initiating the DDR independently of any other factor. SIRT6 has multiple functions in the context of chromatin (Kugel and Mostoslavsky, 2014), including transcriptional regulation. Thus, it might seem somewhat paradoxical that it can initiate the DDR response by merely binding to damage sites. It is not particularly clear how SIRT6 can selectively activate the DDR when bound to DNA damage sites but not when bound to sites of transcription regulation. A possible explanation could rely on the fact that transcription factors are very dynamic, and they usually bind chromatin transiently (Hager et al., 2009). Therefore, we speculate that SIRT6, similarly to MRE11, probes the DNA transiently and even though it is constantly present in chromatin, its binding is not as tight as when encountering broken DNA (as seen in the binding assays) (Myler et al., 2017). Tighter binding of SIRT6 might allow stabilization through protein interactions and modifications, analogous to the processes occurring with MRE11, NBS1, ATM and other DDR proteins. It is also possible that SIRT6 undergoes a conformational change when bound to broken DNA. However, our tethering system suggests that its mere presence in chromatin (in the absence of broken DNA to bind) is sufficient to initiate the DDR cascade.

Interestingly, unlike NBS1, Ku80 and MRE11, SIRT6 recruitment and kinetics are not affected by PARP activity, making it completely independent of parylation and giving it the advantage over other factors that first require parylation for fast recruitment. This feature could be relevant as an adjuvant in cancer treatment (Beck et al., 2014; Haince et al., 2008).

It is also important to note that while SIRT6 can recruit proteins of both HR and NHEJ, and its deficiency affects both pathways, SIRT6 KO impaired the recruitment of MRE11 to sites of laser induced damage. Still, the recruitment of Ku80 was not impaired and in the absence of Ku80, the accumulation of SIRT6 was greater at the sites of damage. It is possible that Ku complex does not require SIRT6 for recognition. Yet, it may require SIRT6 chromatin remodeling activity in later repair steps, while in the initial steps, they may compete for DSB binding.

Our findings place SIRT6 at the top of DSB recognition and possibly as a DDR pathway choice facilitator. Nevertheless, since there is significant cross-talk between the pathways (seen, for example, by the involvement of the HR initiator MRE11 in NHEJ (Xie et al., 2009), it is possible that SIRT6 is involved in both pathways, but plays different roles.

In conclusion, we have demonstrated that Sirtuins and mainly SIRT6 have a role as independent DNA damage sensors, which is critical for the initiation of the DSB-DNA damage response and thereby to support genomic stability and health.

## Acknowledgments

This work was supported by **ISF 188/17** and by the High-tech, Bio-tech and Chemo-tech scholarship of Kreitman School of Advanced Research of Ben Gurion University. We kindly appreciate the plasmids donated by Prof. Misteli and the U20S-LacO cells from Prof. Greenberg. We thank the staff scientist of beamline BM29 of ESRF (Grenoble, France) for providing support and the Israeli Block Allocation Group (BAG) for providing access. We thank Dr. Mario Lebendiker and Dr. Hadar Amartely from the Protein Purification Facility Wolfson Centre for Applied Structural Biology - The Hebrew University of Jerusalem, for their help with the SEC MALS experiments. We thank Dr. Eyal Gur and Dr. Maayan Korman from Ben-Gurion University for their contribution to the development of the NAD^+^ consumption assay. We thank Dr. Amir Aharoni and Dr. Adi Hendler from Ben-Gurion University for their help and advice.

## Supplementary Material and Methods

### Cell cultures

All cells were cultured in DMEM, 4.5g/l glucose, supplemented with 10% fetal bovine serum, 1% Penicillin and Streptomycin cocktail and 1% L-glutamine. Cells were cultured with 5% CO_2_ at 37°C.

### Plasmids and transfections

To prepare pQCXIP-msirt6-GFP-LacR, mouse *sirt6* without stop codon was amplified by PCR and introduced in frame with GFP-LacR into AgeI site of plasmid pQCXIP-GFP-LacR (Addgene, 59418).

pQCXIP-mSIRT6-H133Y-GFP-LacR was prepared by Quick Change Site Directed Mutagenesis of msirt6 flanked by AgeI sites in pGEM, and after sequencing, introduced to AgeI site in frame with the fused GFP-LacR of pQCXIP-GFP-LacR (Addgene, 59418).

pQCXIP-Cherry-LacR was prepared by excision of AgeI / XhoI GFP fragment of pQCXIP-GFP-LacR and exchanged with AgeI / XhoI mCherry amplified from pDEST-mCherry-LacR-BRCA1 (Addgene, 71115).

pQCXIP-mSIRT6-Cherry-LacR was prepared by introducing the AgeI msirt6 from pQCXIP-KU80-GFP-LacR was prepared by introducing KU80, amplified from pEGFP-C1-FLAG-Ku80 (Addgene, 46958) into AgeI site of pQCXIP-GFP-LacR, in fame with GFP.

pQCXIP-hSIRT1-GFP-LacR was prepared by inserting the amplified SIRT1 from SIRT1-Flag (Mostoslavsky Lab) with AgeI site in frame to GFP-LacR of plasmid pQCXIP-GFP-LacR (Addgene, 59418).

pQCXIP-hSIRT2-GFP-LacR was prepared by inserting the amplified SIRT2 from SIRT2-Flag (Addgen #13813) with AgeI site in frame to GFP-LacR of plasmid pQCXIP-GFP-LacR (Addgene, 59418).

pQCXIP-hSIRT7-GFP-LacR was prepared by inserting the amplified SIRT7 from SIRT7-Flag (Addgen #13818) with AgeI site in frame to GFP-LacR of plasmid pQCXIP-GFP-LacR (Addgene, 59418).

pQCXIP-Core hSIRT6-GFP-LacR was prepared by inserting the amplified 233 amino acid core region from amino acid 43 to aa 276 of human SIRT6 and introduced into AgeI site of pQCXIP-GFP-LacR (Addgene, 59418) with additional Methionine before amino acid 43 and in frame to GFP-LacR of plasmid).

pMal-C2-hSIRT6 A13W, D63H, D63Y, W188A, D190Wand I217A were prepared by Quick Change Site-directed Mutagenesis on pMal-C2-hSIRT6. The mutation was affirmed by sequencing.

www.addgene.org/46958/org/46958/P-C1-F

All PCRs were performed with Hot start, KAPA HiFi #KM 2605 or abm Kodaq #G497-Dye proofreading polymerases.

All clones were sequenced for validation, and expression of the fluorescent fusion proteins were checked by transfection into cells.

All transfections were performed using PolyJet^TM^ In Vitro Transfection (SignaGen, SL100688) according to manufacturer’s instructions.

### Immunofluorescence

U2OS cells were washed with PBS and fixed with 2% paraformaldehyde for 15 min at room temperature, followed by ad additional wash. Quenching was then performed with 100mM Glycine for 5 min at RT as well. Cells were permeabilized (0.1% NaCitrate, 0.1% Trition X-100, pH 6, in DDW) for 5 min and washed again. After 1h blocking (0.5% BSA, 0.1% Tween-20 in PBS), cells were incubated with primary antibody diluted in blocking buffer over night at 4°C. The next day, cells were washed three times with wash buffer (0.25% BSA, 0.1% Tween-20 in PBS), incubated for 1h with secondary antibody (diluted in blocking buffer 1:200) at RT and washed three more times. Cells were then DAPI stained for three minutes at RT and washed with PBS twice before imaging.

### Tethering assay

U2OS cell containing 256X LacO sequence repeats in their genome were transfected with plasmids of chimeric LacR-DDR enzyme-GFP/Cherry proteins. Cells were either co-transfected with a second plasmid of a fluorescent/ Flag tagged protein or Immuno-stained (See Immunofluorescence) for an endogenic protein. Cells exhibiting visible foci of LacR-DDR-GFP/Cherry at LacO sites were located using fluorescent microscopy, while co-localization between both proteins was assessed visually using Olympus CellSens Software. Positive co-localization percentage was calculated and compared to co-localization percentage with LacR-GFP/Cherry.

• Notes:

pQCXIP-Ku80-GFP-LacR plasmid used in this assay contains Ku80 that was acquired from Addgene (cat. #46958) and contains D158G mutation.

pQCXIP-SIRT1-GFP-LacR plasmid used in this assay contains SIRT1 that was obtained from Mostoslavsky lab (Zhong et al., 2010). This protein variant is lacking 79 amino acids in the N-terminus.

### Immunoprecipitation (IP)

Flag tagged proteins were purified form transfected HEK293T cells. Cells were collected and washed with PBS. Cell disruption was performed in Lysis buffer (0.5M KCl, 50mM Tris-HCl pH7.5, 1% NP40, 0.5M DTT, 200mM TSA and protease and phosphatase inhibitors in DDW) by 10 min rotation at 4°C. Cell debris were sedimentated by 15 min centrifugation at 21,000g. Lysate was collected and added to ANTI-FLAG M2 Affinity Gel (SIGMA-ALDRICH, A2220) beads for 2 hr rotation at 4°C. Beads were then washed three times with lysis buffer and once with SDAC buffer (50mM Tris-HCl pH9, 4mM MgCl, 50mM NaCl, 0.5mM DTT, 200mM TSA and protease and phosphatase inhibitors in DDW). Proteins were released by flag-peptide.

### Expression and purification of recombinant SIRT6 in *E.coli*

Expression and purification of His-tagged and MBP-tagged proteins in *E.coli* were performed as previously described at Gertman et al., 2018.

### Fluorescence Recovery after Photobleaching (FRAP)

FRAP experiments (Laser induced damage) were performed as previously described at Toiber et al., 2013. In brief, cells were plated in Ibidi µ-Slide 8 well glass bottom plates (Cat.No: 80827) and transfected with the desired fluorescent plasmid. Pre-sensitization with Hoechst (1mM) was done for 10 min before the experiment. FRAP experiments were carried out using a Leica SP5 microscope (German Cancer Research Center (DKFZ) and BioQuant, Heidelberg, Germany) or using a LSM880 microscope (Ben Gurion University, Be’er Sheva, Israel) with a 63X oil immersion objective. Images were acquired in 512 × 512 format with a scan speed of 1,400 Hz. Circular bleach spots with 2 µm diameter were used, which were positioned either at a damage site or at a distant reference site. Spots were bleached with an argon laser at 488 nm with a power of 1 mW in the back aperture of the objective. Images were taken in 5 or 3 seconds intervals, with 3 baseline images taken before bleaching. Acquisition of the bleaching was used for normalization of each cell intensity (average of the baseline intensity of the whole cell nucleus prior to DNA damage). Images analysis was performed using ImageJ 1.52i software.

In Fig. 1A-B and S1A-B, U2OS Cells were incubated with 10uM BrdU or 10uM BrdU + 10uM Olaparib over-night, before going through the same procedure.

### DNA binding assay

Open ended plasmids were prepared in advance by incubating DR-GFP plasmid with EcoRV for blunt ends, KpnI for 3’ over hang or SalI for 5’ over hang according to manufacture instructions. Circular plasmids were subjected to the same conditions with no restriction enzyme.

To achieve protein-DNA binding, flag tagged proteins that were previously immunoprecipitated were incubated at 37°C for 1h with same amount of circular or open ended DNA, 1:5 of 5X Deacetylation buffer (50mM Tris HCl pH8, 50mM NaCl, 4mM MgCl_2_ and 0.5mM DTT in DDW), 1:50 50X Protease inhibitors in DDW.

ANTI-FLAG M2 Affinity Gel (SIGMA-ALDRICH, A2220) beads were blocked with 5% BSA supplemented to 1X Deacetylation buffer (with 1% Phosphates inhibitors) by rotation of 1h in 4°C. Beads were then centrifuged (1000g, 3min, 4°C) and buffer was changed to clean Deac. Buffer 1X. Beads were then distributed equally between all samples.

To achieve beads-protein binding, samples were rotated for 2h in 4°C. After rotation, samples were centrifuged (1000g, 3min, 4°C) and washed 3 times with 1ml of wash buffer (0.1% SDS, 0.5% Triton x-100, 2mM EDTA, 20mM Tris-HCl pH8 and 150mM NaCl in DDW).

Protein-DNA complexes were then released by 2 times 20 min vortex at room temperature with 100ul Elution buffer (0.1M NaHCO_3_ and 1% SDS in DDW).

For His-tagged proteins (acquired from PROSPEC) the assay was performed using HisPur Ni-NTA Resin (ThermoFisher, 88221) at the same conditions with the appropriate buffers (Binding buffer: 20mM Tris HCl pH8, 150mM NaCl, 10% PMSF, 1% phosphatase inhibitors; Wash Buffer: 20mM Tris HCl pH8, 150mM NaCl, 20mM Imidazole; Elution buffer: 20mM Tris HCl pH8, 150mM NaCl, 500mM Imidazole).

• Notes:

SIRT1-Flag used in this assay was obtained from Mostoslavsky lab (PDMI: 20141841). This protein variant is lacking 79 amino acids in the N-terminus.

SIRT1-His used in this assay was acquired from PROSPEC (https://www.prospecbio.com/sirt1_human). This SIRT1 is a 280 amino acids polypeptide (a.a. 254-495).

### DNA isolation

To the eluted DNA from the DNA binding assay 1:1 volume of phenol: chloroform: isoamyl alcohol (25:24:1) was added, vortexed and centrifuged at room temperature for 5 min at 17,000g. Top aqueous layer was then isolated and washed with 1 volume of chloroform: isoamyl alcohol (24:1). Samples were then centrifuged at same conditions, and top aqueous layer was isolated. 1/10 volume 3M NaOAc, 30ug glycogen and 2.5 volumes ice cold 100% EtOH were added to each sample, followed by an incubation for at least 30min at −80°C. After incubation, DNA was precipitated by centrifugetion at max speed for 30min, 4°C, supernatant was discarded and the pellet washed with 500ul 70% ice-cold EtOH. Samples were then centrifuged at max speed for 30min, 4°C, supernatant was discarded and DNA pellet was air dried before re-suspension with ultra-pure water.

### Quantitative PCR

For relative quantification of the DNA isolated from the all DNA binding assays performed, qPCR was performed using SsoAdvanced^TM^ Universal SYBER^®^ Green Supermix (BIO-RAD, 172-5274) according to manufacturer’s instructions.

Primers used for DRGFP plasmid amplification:

Forward: 5’-TCTTCTTCAAGGACGACGACGGCAACT-3’ Reverse: 5’-TTGTAGTTGTACTCCAGCTTGTGC-3’

### Exonuclease assay

DR-GFP plasmid was cut with restriction enzymes generating linear DNA with blunt (EcoRV) or overhanging ssDNA (SalI or KpnI). DNA cleavage was confirmed by agarose gel electrophoresis. 10ug of the restricted DNA was incubated with BSA, NBS1, MRE11 or SIRT6 purified proteins in NEB exonuclease buffer for 0 to 20 min. ExoI was then added to the samples. Samples of each reaction were taken at 0, 10 and 20 min.

DNA was purified by Qiagen PCR purification kit. The purified DNA was run on 0.8% agarose Gel, and DNA amount was assessed by image analysis using ImageJ 1.52i software and normalized to the amount of the DNA at 0’ time point.

### Fluor de lys (FDL) activity assay

Fluor de lys assay with SIRT6-point mutant-MBP proteins was performed as previously described at Gertman et al., 2018.

### NAD^+^ consumption assay

Purified SIRT6-Flag was incubated at 37°C for 3h with either PstI digested pDR-GFP (DSB), ssDNA or a H3K56 acetylated peptide with 2.5mM NAD^+^ and HEPES buffer (50mM HEPES pH 7.5, 100mM KCl, 20mM MgCl_2_, 10% Glycerol). After incubation samples were supplemented with 1uM 1,3-Propanediol dehydrogenase (1,3-PD) and 170mM 1,3 Propanediol for an additional 3h incubation. NAD^+^ consumption by SIRT6 was assessed by NADH levels produced by 1,3-PDase activity, by measuring its absorption at 340nm. To monitor spontaneous NAD^+^ consumption in the presence of PstI digested pDR-GFP, ssDNA or H3K56 acetylated peptide, the assay was conducted without SIRT6, and each treatment was normalized to its control.

### EMSA

SIRT6 in storage buffer (20mM Tris-HCl, pH 7.4; 150mM NaCl; 50% glycerol) was equilibrated with DNA (or RNA) for 20 minutes on ice. The buffer composition of EMSA was optimized to obtain the maximum resolution for resolving DNA/RNA. Reactions (final volume 10μL) were resolved by electrophoresis at 4°C through native gel containing 5% (for blunt-end and sticky-end DNA), 8% (for ssDNA) and 10% (for RNA) polyacrylamide (29:1 acrylamide: bisacrylamide) in 1X TBE buffer. Autoradiographs of the dried gels were analyzed by densitometry using Fujifilm PhosphorImager. The signal was quantified by ImageQuant TL.

GraphPad Prism 7 was used to estimate apparent Kd value for ssDNA (one site, specific binding fit, y = B_max_[SIRT6]/(K_d_ + [SIRT6])) and for blunt-end and sticky-end DNA (specific binding with Hill slope, y = B_max_[SIRT6]^h^/(K_d_ ^h^ + [SIRT6]^h^)).

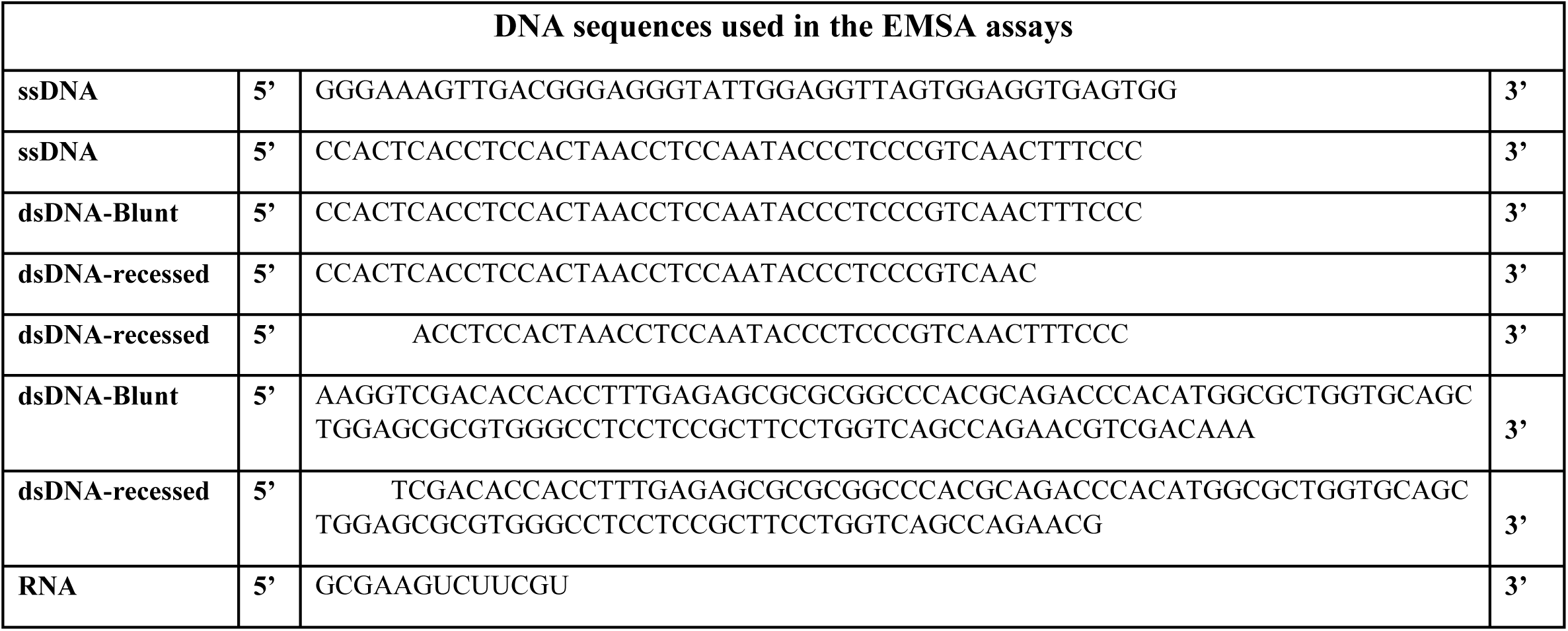

### SAXS

SAXS data were collected at BioSAXS beamline BM29 (ESRF, Grenoble, France), possessing Pilatus 1M detector. The scattering intensity was recorded in the interval 0.0035 < q < 0.49 Å^−1^. The measurements were performed at 20 °C. SIRT6 (alone or in the presence of dsDNA) was measured at concentration of 0.5 mg/ml, as it tends to aggregate at higher concentrations. The scattering of the buffer was also measured and subtracted from the scattering of the samples by using Primus (P.V. Konarev et al.; 2003; J Appl Cryst. 36, 1277-1282.).

PyMOL (https://pymol.org/) was used to extract the structures of SIRT6 dimer and tetramer from the available crystal structure (PDB ID code: 3pki). CRYSOL (Svergun DI et al.; 1995; J Appl Crystallogr 28:768–773) was then used to compute the artificial SAXS spectra of each protein specie. These spectra served as a reference for the reconstitution of experimental SAXS data:

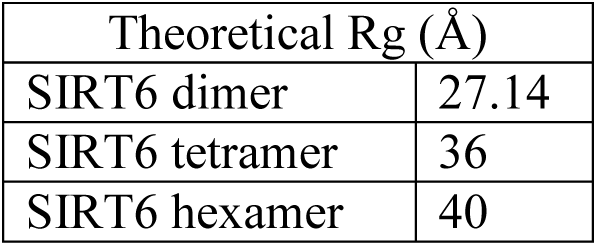

Values for the radius of gyration (R_g_) and maximum particle dimension (D_max_) were derived from distance distribution function *P(r)*, using in-house script (Akabayov B. et al.; 2010; PNAS; 107(34):15033*–*15038). This script was designed to perform an automatic search for the best fitting parameters in GNOM (Svergun DI; 1992; J Appl Crystallogr 25:495–503).

In the end, DAMMIN (Svergun DI; 1999; *Biophys. J*., 2879-2886) was used to reconstruct the molecular envelope based on the best GNOM fit (obtained from the script analysis and refined manually). 18 models were calculated and averaged using DAMMAVER (Svergun DI; 2003; *J. Appl. Cryst. 36:860-864.*).

### SEC-MALS

A miniDAWN TREOS multi-angle light scattering detector, with three angles (43.6°, 90° and 136.4°) detectors and a 658.9 nm laser beam, (Wyatt Technology, Santa Barbara, CA) with a Wyatt QELS dynamic light scattering module for determination of hydrodynamic radius and an Optilab T-rEX refractometer (Wyatt Technology) were used in-line with a size exclusion chromatography analytical column, Superdex 200 Increase 10/300 GL (GE, Life Science, Marlborough, MA) equilibrated in buffer (50mM tris, 150 mM NaCl and 4 mM MgCl2, pH 8.0).

Experiments were performed using an AKTA explorer system with a UV-900 detector (GE), at 0.8ml/min. All experiments were performed at room temperature (25 °C). Data collection and mass calculation by SEC-MALS analysis were performed with ASTRA 6.1 software (Wyatt Technology). The refractive index of the solvent was defined as 1.331 and the viscosity was defined as 0.8945 cP (common parameters for PBS buffer at 658.9 nm). dn/dc (refractive index increment) value for all samples was defined as 0.185 mL/g (a standard value for proteins). For SIRT6 experiment-150ul 4.5mg/ml human-SIRT6-His was injected. For SIRT6+DNA-200ul human-SIRT6-His + 50ul DNA after 1 h incubation at 37°C was injected.

### Statistical analysis

Statistical analysis was done using GraphPad Prism 7. Analysis included either one-way or two-way ANOVA followed by a post-hoc Dunnet test or Tukey test respectively. Significance was set at p< 0.05.

For all DNA binding assay results, statistical analysis was predeceased by logarithmic transformation to overcome large variance between the different experiments. Statistical analysis was performed on the transformed data as described.

---

**Supplementary Table 1:** Plasmids.

| **Plasmid** | **Source** | **Cat. number** | **PMID** |
| --- | --- | --- | --- |
| ATM-LacR-Cherry | Misteli Lab | - | 18483401 |
| CDT1-TagRFP | ThermoFisher | P36237 | 18267078 |
| CMV-Flag | Mostoslavsky Lab | - | 23911928 |
| MRE11-Flag | Mostoslavsky Lab | - | 18678890 |
| MRE11-LacR-Cherry | Misteli Lab | - | 18483401 |
| mRFP-SIRT6 | Jackson Lab | - | 20829486 |
| NBS1-Flag | Mostoslavsky Lab | - | 18678890 |
| NBS1-LacR-Cherry | Misteli Lab | - | 18483401 |
| pcDNA3.1(+)Flag-His-ATM-WT | Addgene | 31985 | 9733515 |
| pcDNA5-FRT/T0-Flag-53BP1 | Addgene | 52507 | 23333306 |
| pDEST 3x Flag-pcDNA5-FRT/T0-BRCA1 | Addgene | 52504 | 23333306 |
| pEGFP-C1-FLAG-Ku70 | Addgene | 46957 | 23897892 |
| pEGFP-C1-FLAG-Ku80 | Addgene | 46958 | 23897892 |
| pEGFP-C1-FLAG-XRCC4 | Addgene | 46959 | 23897892 |
| pEGFP- SIRT6 | Jackson Lab | - | 20829486 |
| pET28 hSIRT6-His | Aharoni Lab | - | 29476161 |
| pHPRT-DRGFP | Addgene | 26476 | 11751629 |
| pMal-C2-hSIRT6-WT | Aharoni Lab | - | 29476161 |
| pMal-C2-hSIRT6-A13W | Toiber Lab | - | - |
| pMal-C2-hSIRT6-D63H | Toiber Lab | - | - |
| pMal-C2-hSIRT6-D63Y | Toiber Lab | - | - |
| pMal-C2-hSIRT6-H133Y | Aharoni Lab | - | 29476161 |
| pMal-C2-hSIRT6-D188A | Toiber Lab | - | - |
| pMal-C2-hSIRT6-D190W | Toiber Lab | - | - |
| pMal-C2-hSIRT6-I217A | Toiber Lab | - | - |
| pQCXIP-Cherry-LacR | Toiber Lab | - | - |
| pQCXIP-SIRT6 Core-GFP-LacR | Toiber Lab | - | - |
| pQCXIP-GFP-LacR | Addgene | 59418 | 23929981 |
| pQCXIP-KU80-GFP-LacR | Toiber Lab | - | - |
| pQCXIP-mSIRT6-Cherry-LacR | Toiber Lab | - | - |
| pQCXIP-mSIRT6-GFP-LacR | Toiber Lab | - | - |
| pQCXIP-mSIRT6-H133Y-GFP-LacR | Toiber Lab | - | - |
| pQCXIP-SIRT1-GFP-LacR | Toiber Lab | - | - |
| pQCXIP-SIRT2- GFP-LacR | Toiber Lab | - | - |
| pQCXIP-SIRT7- GFP-LacR | Toiber Lab | - | - |
| SIRT1-Flag | Mostoslavsky Lab | - | 20141841 |
| SIRT2-Flag | Addgene | 13813 | 12620231 |
| SIRT3-Flag | Addgene | 13814 | 12620231 |
| SIRT4-Flag | Addgene | 13815 | 12620231 |
| SIRT5-Flag | Addgene | 13816 | 12620231 |
| SIRT6 Core | Chua Lab | - | 20117128 |
| SIRT6-WT-Flag | Mostoslavsky Lab | - | 20141841 |
| SIRT7-Flag | Addgene | 13818 | 12620231 |
| SNF2H-WT-GFP-LacR | Goodarzi Lab | - | 25533843 |

**Supplementary Table 2:** Antibodies.

| Antibody | Company | Cat. num' | Method | Dilution |
| --- | --- | --- | --- | --- |
| Alexa Fluor® 488 AffiniPure Donkey Anti-Rabbit IgG (H+L) | Jackson Immunoresearch | 711-545-152 | IF | 1:200 |
| Alexa Fluor® 594 AffiniPure Donkey Anti-Rabbit IgG (H+L) | Jackson Immunoresearch | 711-585-152 | IF | 1:200 |
| Alexa Fluor® 488 AffiniPure Donkey Anti-Mouse IgG (H+L) | Jackson Immunoresearch | 715-545-150 | IF | 1:200 |
| Alexa Fluor® 594 AffiniPure Goat Anti-Mouse IgG (H+L) | Jackson Immunoresearch | 115-585-062 | IF | 1:200 |
| 53BP1 | Snata-Cruz Biotechnology | sc-22760 | IF | 1:300 |
| BRCA1 | Snata-Cruz Biotechnology | sc-7298 | WB | 1:1000 |
| Flag | Sigma-Aldrich | F1804 | IF/WB | 1:900/1:1000 |
| Flag Beads | Sigma-Aldrich | A2220 | IP | - |
| gamma H2A.X (phospho s139) | abcam | ab2893 | IF/WB | 1:3000/ 1:1000 |
| Goat Anti-Rabbit IgG H&L (HRP) | abcam | ab6721 | WB | 1:10000 |
| Histone H3 | abcam | ab1791 | WB | 1:5000 |
| Histone H3 (acetyl K56) | abcam | ab76307 | WB | 1:1000 |
| HSC 70 | Snata-Cruz Biotechnology | sc-7298 | WB | 1:1000 |
| Ku80 | Cell Signaling | #2180 | WB | 1:1000 |
| MRE11 | abcam | ab214 | WB | 1:1000 |
| phospho-ATM (Ser1981) | Cell Signaling | #5883 | WB | 1:1000 |
| phospho-Histone H2A.X (Ser139) | Millipore | 05-636 | IF | 1:1500 |
| Rabbit Anti-Mouse IgG H&L (HRP) | abcam | ab97046 | WB | 1:10000 |
| RNF8 | Snata-Cruz Biotechnology | Sc-271462 | WB | 1:1000 |
| SIRT6 | abcam | ab62739 | WB | 1:1000 |
| Tubulin | Merck | MAB1637 | WB | 1:1000 |
| Ku80 | Cell Signaling | #2180 | WB | 1:1000 |

**Supplementary Table 3:** Proteins-DNA binding assay.

| Protein | Company | Cat. num' |
| --- | --- | --- |
| SIRT1 Human | PROSPEC | PRO-1909 |
| SIRT3 Human | PROSPEC | PRO-462 |
| SIRT5 Human | PROSPEC | PRO-1774 |
| SIRT6 Human | PROSPEC | PRO-282 |

**Supplementary Figure 1.**
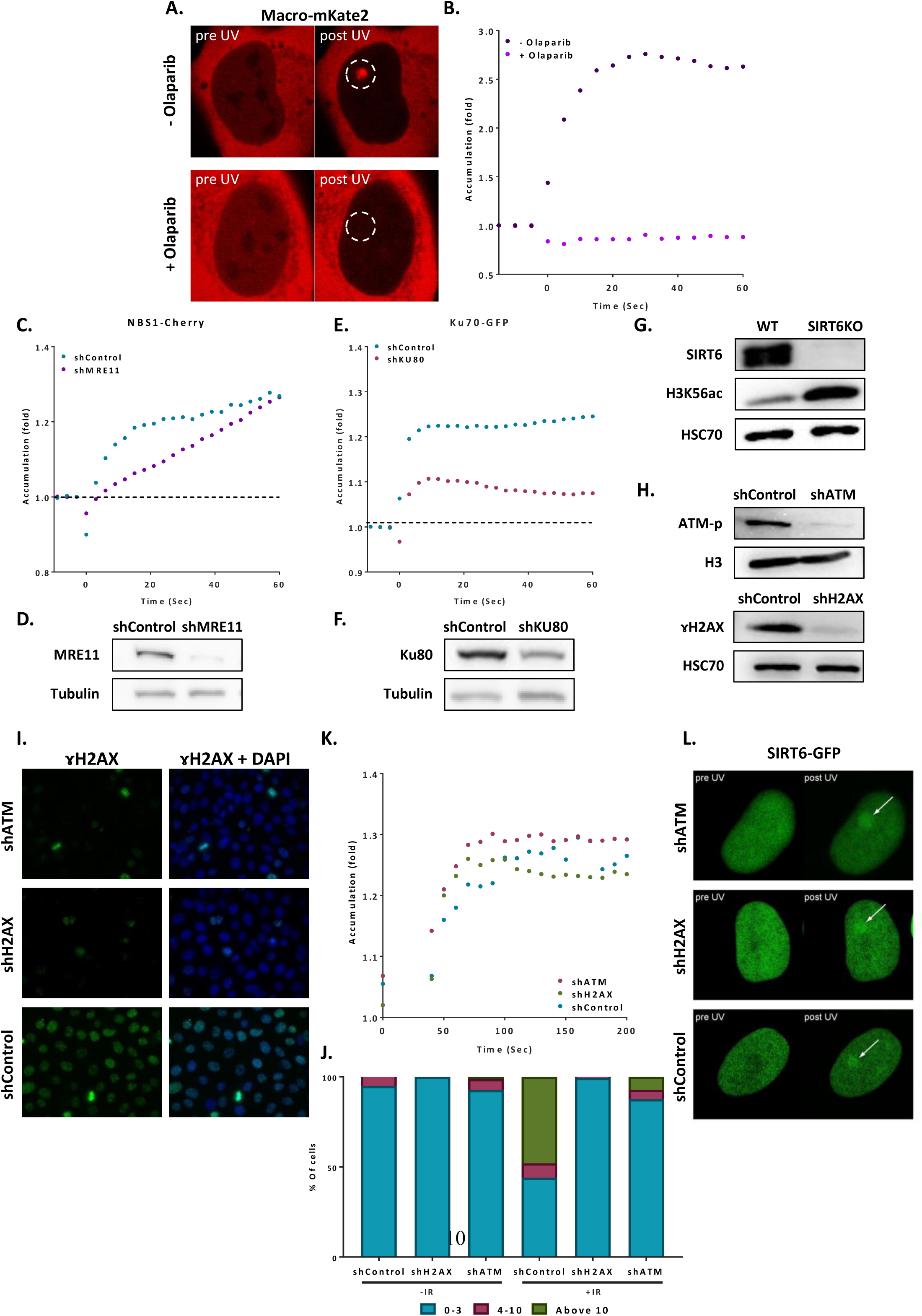
**(A-B)** Live imaging for Macro-domain mKate2 upon laser-induced damage (LID) in naïve U2OS cells treated with BrdU and Olaparib (n=10) and cells treated with BrdU alone (n=10). Graph presents accumulation of the Macro-domain over time, measured by change in fluorescence at 5sec intervals. **(C)** Accumulation over time of NBS1-LacR-Cherry at sites of LID in control Hela cells (n=10) or in cells knocked-down for MRE11 (n=15), both treated with Hoechst, measured by change in fluorescence at 3sec intervals. **(D)** Protein blot of nuclear extraction of control and shMRE11 Hela cells. **(E)** Accumulation of KU70-Flag-GFP at sites of LID in control (n=15) Hela cells or in cells knocked-down for KU80 (n=15), both treated with Hoechst, measured by change in fluorescence at 3sec intervals. **(F)** Blot of control and shKU80 Hela cells. **(G)** Blot of SIRT6 KO and WT U2OS cells. **(H)** Blot of nuclear protein extraction of Hela cells knocked-down for ATM, H2AX or RNF8 and control cells. **(I)** IF of ɤH2AX in sh-Hela cells upon irradiation. **(J)** ɤH2AX foci quantification in Control (n[-IR]=283, n[+IR]=301), shH2AX (n[-IR]=284, n[+IR]=322) and shATM (n[-IR]=299, n[+IR]=385) Hela cells. The graph presents the percentage of cells that present either 0-3, 4-10 or above 10 foci per nucleus, with or without exposure to irradiation. The graph depicts an average of 6 experiments. **(K-L)** Accumulation of SIRT6-GFP at sites of laser induced damage in Hela cells knocked-down for ATM (n=10), H2AX (n=10) or control (n=10). Graph presents accumulation of SIRT6 over time, measured by change in fluorescence.

**Supplementary Figure 2.**
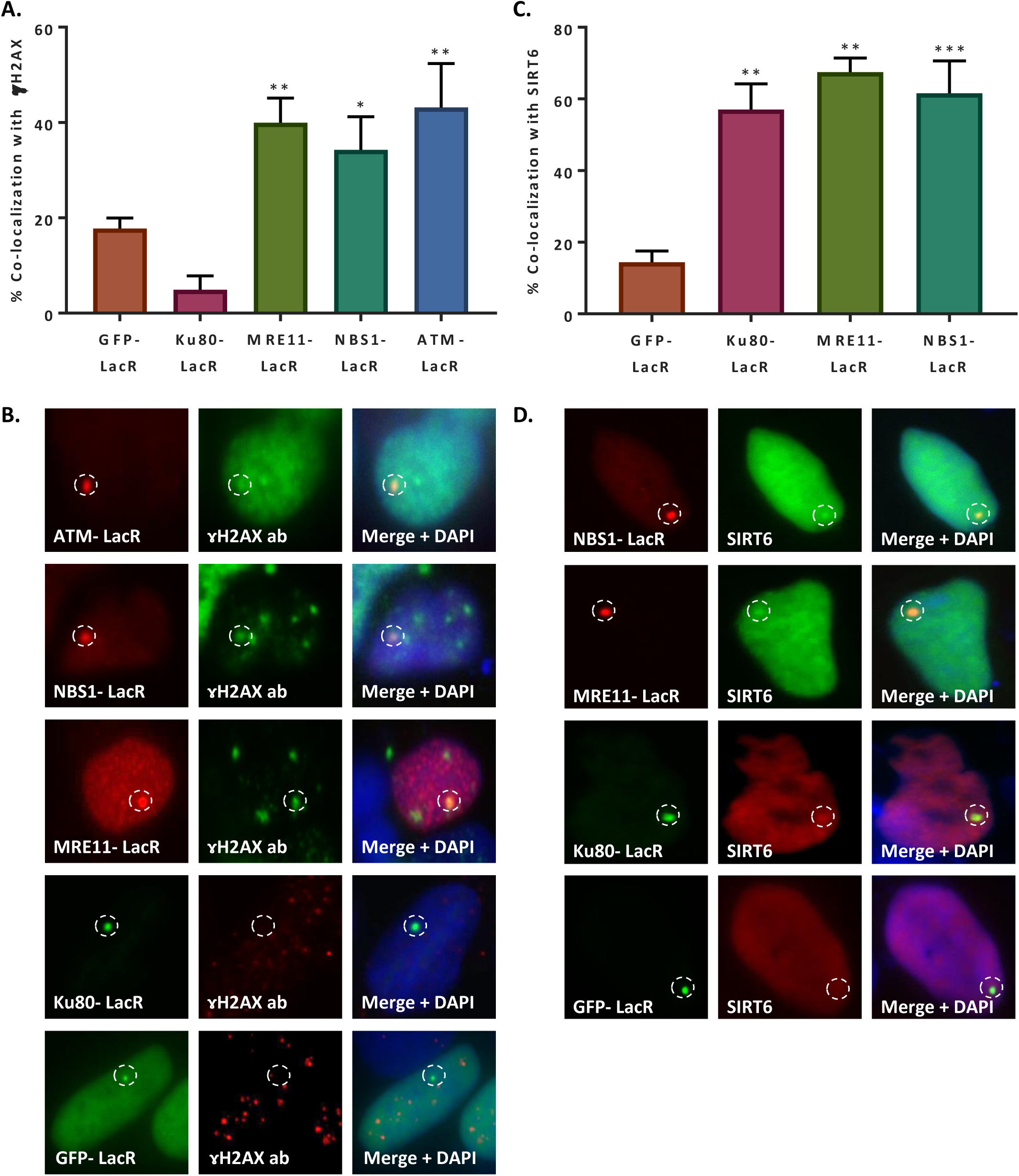
**(A-B)** Initiation of DNA damage response by LacR-ATM Cherry (n=33, p<0.005), LaR-NBS1 Cherry (n=82, p<0.05), LacR-MRE11 Cherry (n=136, p<0.005) and LacR-KU80 GFP (n=52, p>0.05) in the tethering system, shown by IF and calculation of co-localization percentage with ɤH2AX (compared to LacR-GFP (n=310)). **(C-D)** Recruitment of SIRT6-GFP/SIRT6-C1h2erry to LacO sites by LacR-NBS1 Cherry (n=87, p<0.0005), LacR-MRE11 Cherry (n=31, p<0.005), LacR-KU80 GFP (n=45, p<0.005) and LacR-GFP (n=85). The graph depicts averages of 3-5 experiments +/- SEM.

**Supplementary Figure 3.**
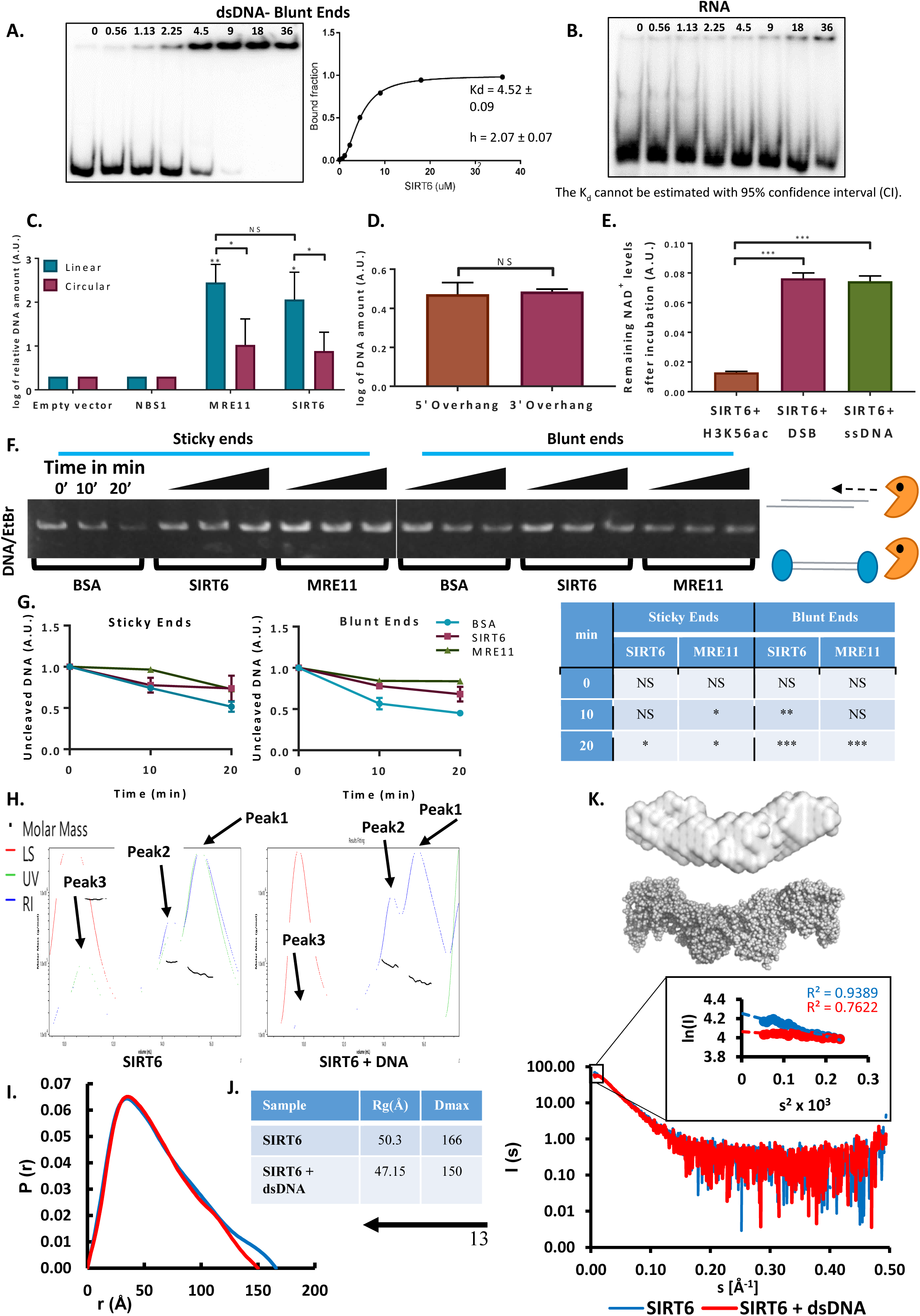
**(A)** Gel retardation assay of 32P-5’ end-labeled dsDNA with blunt ends as a function of increasing concentrations of SIRT6-His. Kd value (Kd=3.59 ± 0.17). **(B)** Gel retardation assay of 32P-5’ end-labeled RNA as a function of increasing concentrations of SIRT6-His. Kd could not have been calculated with 95% confidence interval (CI). **(C)** DNA binding ability of SIRT6-Flag, MRE11-Flag and NBS1-Flag (as a control) of Circular and linear plasmids, assessed by a DNA binding assay. The graph depicts averages of 3-9 experiments +/- SEM, after logarithmic transformation. **(D)** SIRT6-Flag DNA binding ability of plasmids with 3’ over-hang and 5’ over-hang assessed by a DNA binding assay. The graph depicts log of averages of 3 experiments +/- SEM. **(E)** NAD+ consumption assay test whether SIRT6 breaks NAD+ in the presence of DNA alone or an acetylated peptide (H3K56ac). **(F)** Protection of sticky ended and blunt ended plasmids by BSA, SIRT6 and MRE11 against Exonuclease 1 activity in different time points. Averages +/- SEM of 3 experiments. **(G)** Statistical analysis of the End protection. Un-cleaved DNA amounts in the presence of SIRT6 or MRE11 were compared to the amounts measured in the presence of BSA (as a control) at the 3 different time points. (*=p<0.05, **=p<0.005, ***=p<0.0005). **(H)** SEC-MALS analysis of SIRT6 or SIRT6 with dsDNA 10bp oligo with 3 overhanging ends on both sides. **For SIRT6:** <u>Peak 1:</u> Protein mass (calculated by UV) = 66.7±3.3 kDa, Protein mass (calculated by RI) = 70.7±3.5 kDa. <u>Peak2:</u> Protein mass (calculated by UV) = 114.3±5.7 kDa, Protein mass (calculated by RI) = 103.8±4.9 kDa. <u>Peak3:</u> Protein mass (calculated by UV) = 609.7±14.6 kDa, Protein mass (calculated by RI) = 833.6±16.4 kDa. **For SIRT6+DNA:** <u>Peak 1:</u> Protein mass (calculated by UV) = 52.5±3.1 kDa, Protein mass (calculated by RI) = 55.7±3.2 kDa. <u>Peak2:</u> Protein mass (calculated by UV) = 99.4±3.8 kDa, Protein mass (calculated by RI) = 94.6±3.5 kDa. <u>Peak3:</u> Protein mass (calculated by UV) = extremely high, Protein mass (calculated by RI) = 9224±258 kDa. **(I)** X-ray scattering profile (right) and the distance distribution function (left) of SIRT6 (blue) and SIRT6 bound to dsDNA (red). **(J)** Overall parameters for small angle X-ray scattering of SIRT6 alone and SIRT6 bound to dsDNA determined from the distance distribution function P(r). Rg is the radius of gyration and Dmax, maximum dimension of the particle **(K)** SAXS structure of SIRT6 (grey surface). Ab initio models were reconstructed from SAXS data using the computer program DAMMIN [D. I. Svergun (1999) Restoring low resolution structure of biological macromolecules from solution scattering using simulated annealing. Biophys J. 2879-2886] and were averaged by the computer program DAMAVER [V. I. Volkov and D. I. Svergun (2003). Uniqueness of ab-initio shape determination in small-angle scattering. J. Appl. Cryst. 36, 860-864]. The crystal structure of SIRT6 tetramer (grey spheres) was extracted from the crystal structure of SIRT6 (pdb id: 3PKI) and compared with the obtained SAXS model in PyMOL (http://www.pymol.org).

**Supplementary Figure 4.**
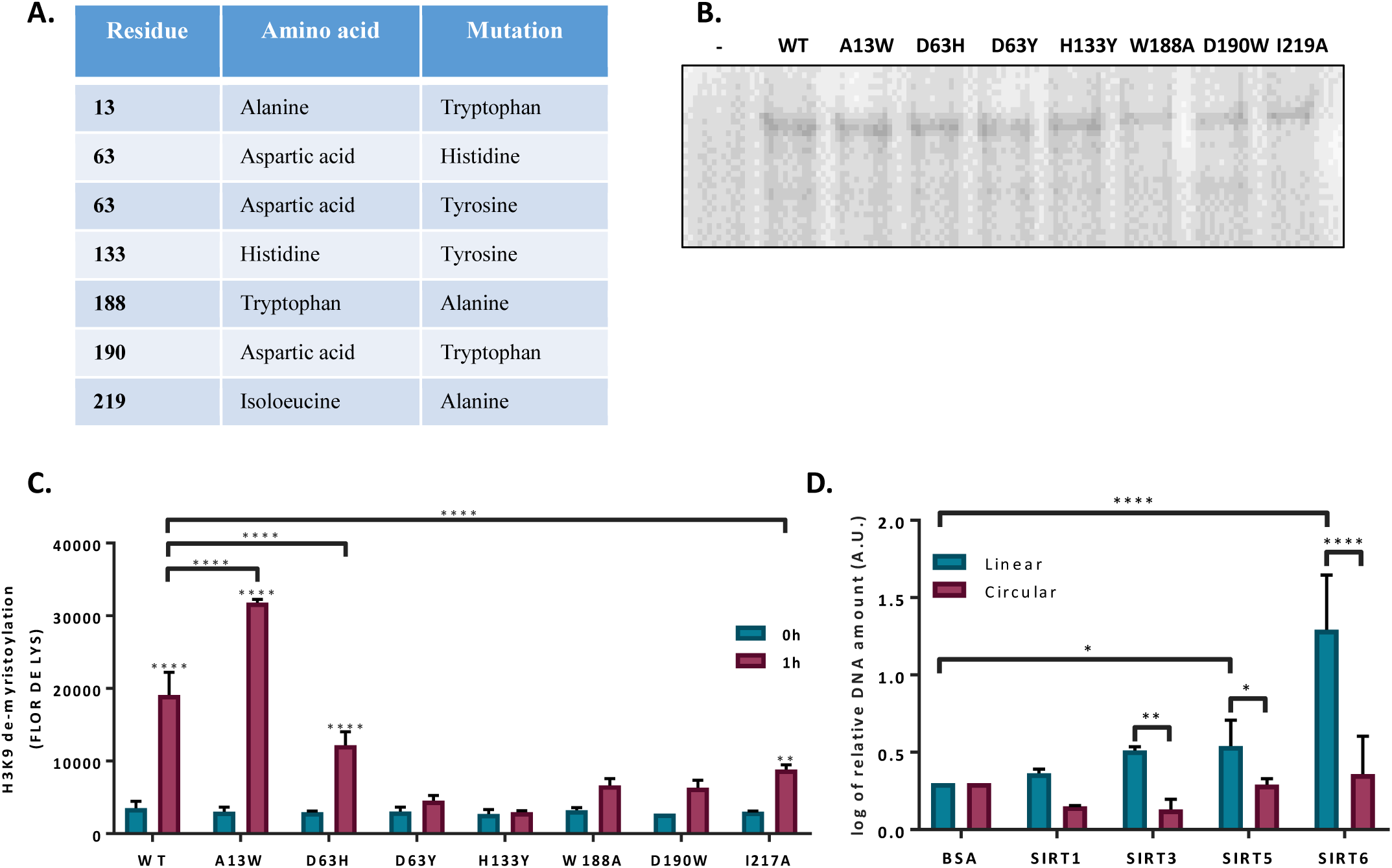
**(A)** List of generated SIRT6 point mutations in the SIRT6 tunnel structure. **(B)** Ponceau staining of SIRT6- MBP pint mutants. **(C)** SIRT6-MBP mutants catalytic activity, assessed by H3K9 de-myristolation in a FLOR DE LYS assay. **(D)** His tagged mammalian Sirtuins DNA binding ability of circular and linear plasmids. The graph depicts log of averages of 4-7 experiments +/- SEM. (*=p<0.05, **=p<0.005, ***=p<0.0005).

**Supplementary Figure 5.**
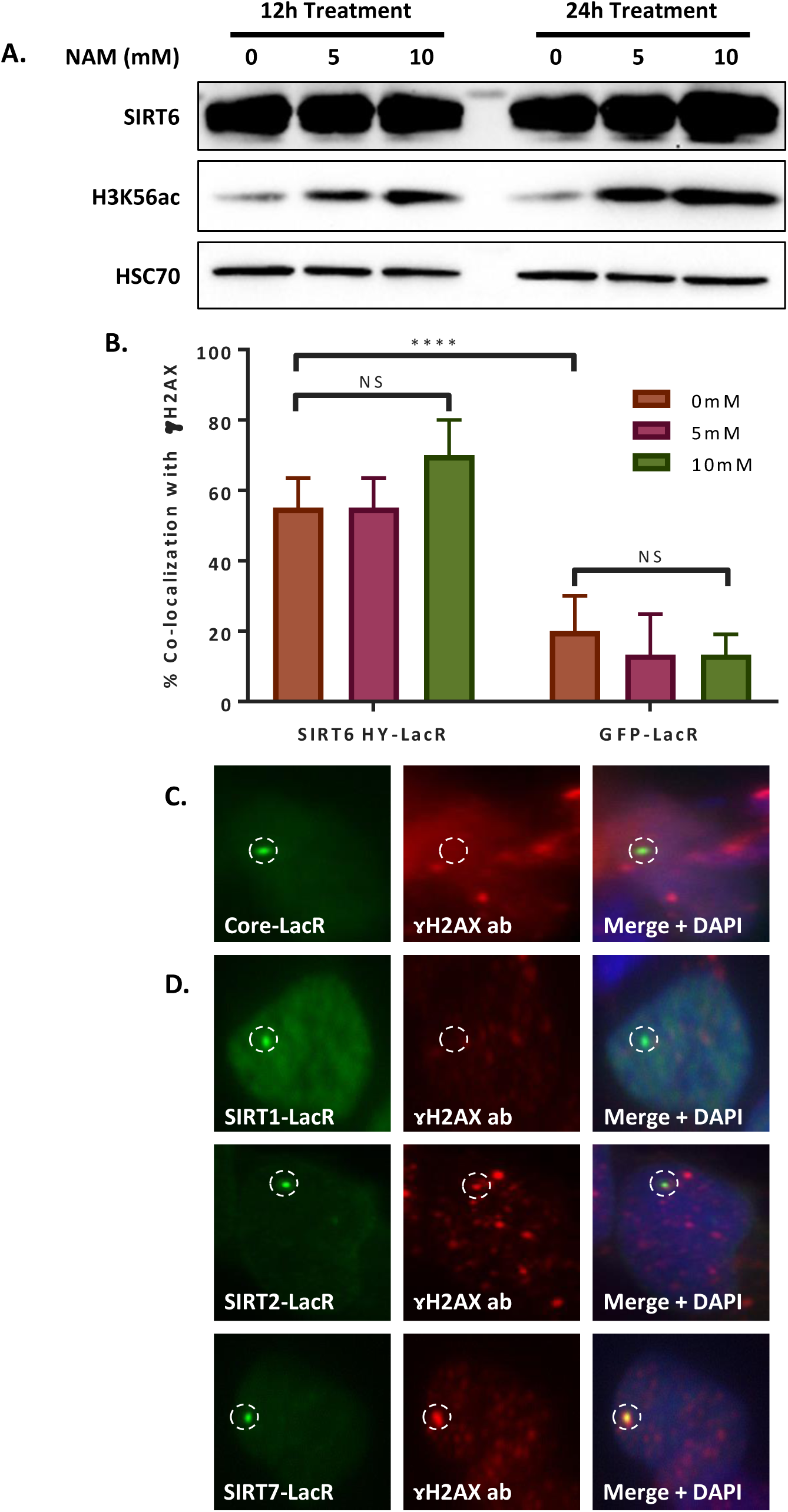
**(A)** Western blot of nuclear protein fraction of LacO containg U2OS cells after 12h or 24h treatment with 0, 5 or 10mM Nicotinamide (NAM) supplemented to their media. **(B)** Co-localization percentage with ɤH2AX measured for LacR-SIRT6-HY GFP and LacR-GFP after 24h treatment with NAM at three different concentrations. (*=p<0.05, **=p<0.005, ***=p<0.0005). **(C)** Images of IF of co-localization of LacR-SIRT6 core domain-GFP (n=66, p>0.05). Experiment quantification appears in Fig. 5C. **(D)** Images of IF of co-localization of LacR-SIRT1 GFP (n=44, p>0.05), LacR-SIRT2 GFP (n=67, p<0.005) and LacR-SIRT7 GFP (n=68, p<0.005). Experiment quantification appears in Fig. 5D.

**Supplementary Figure 6.**
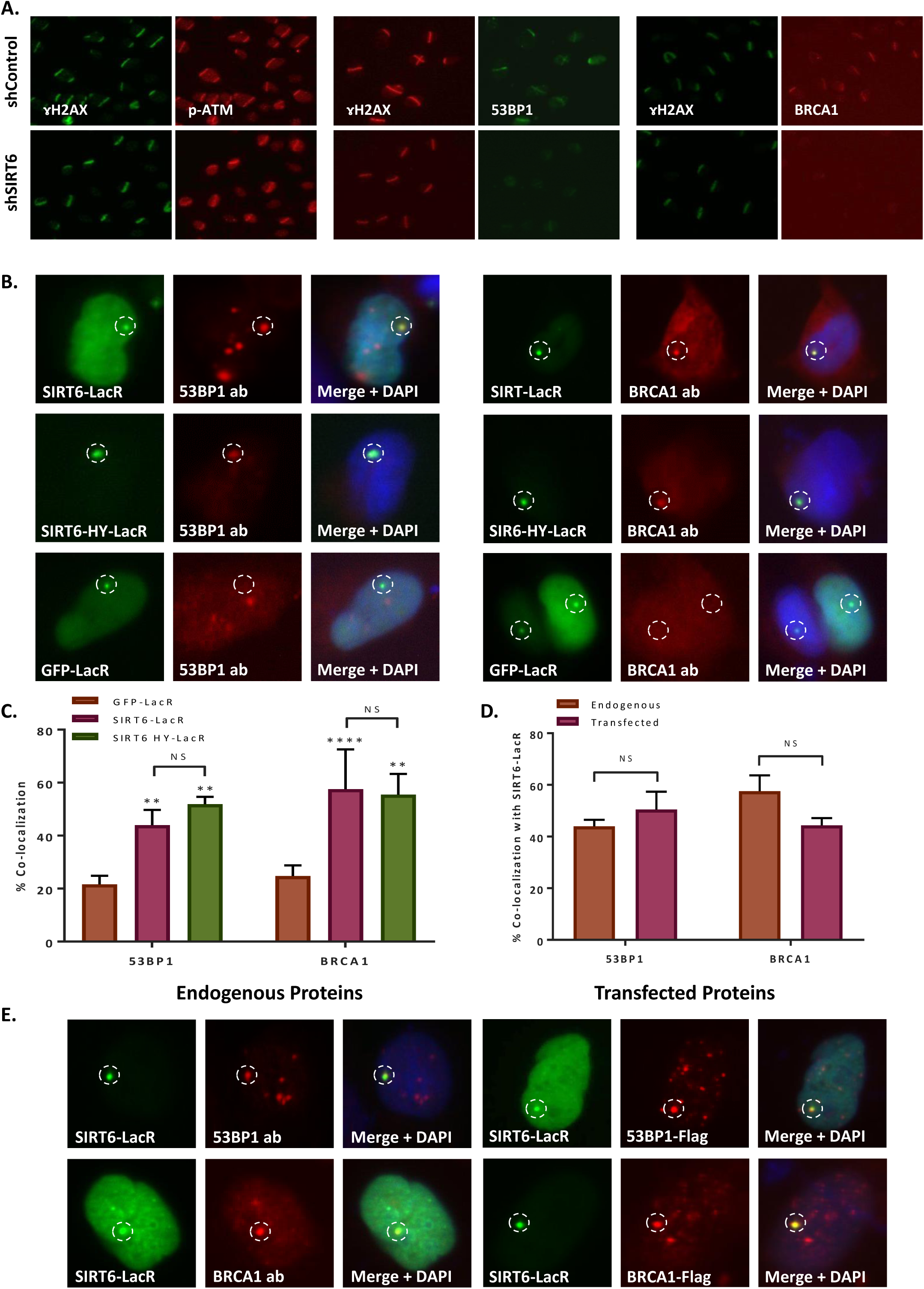
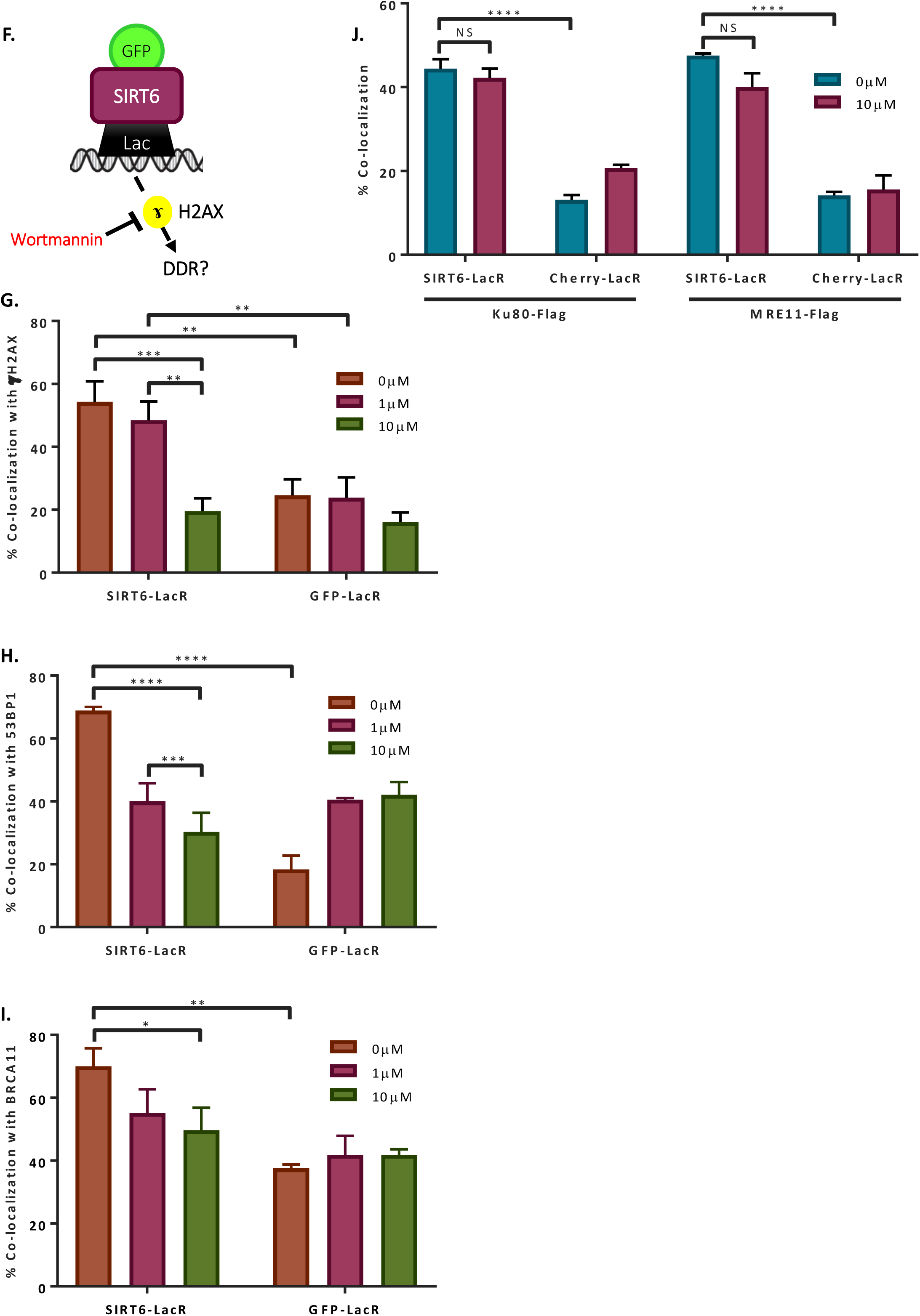
**(A)** Laser induced damaged followed by IF at 30 min after damage. Recruitment of p-ATM, 53BP1 or BRCA1 in U20S shCtrl or shSIRT6. **(B-C)** IF of co-localization of 53BP1 and BRCA1 with LacR-SIRT6 GFP (53BP1: n=247, p<0.005; BRCA1: n=223, p<0.005) and LacR-HY GFP (53BP1: n=113, p<0.005; BRCA1: n=122, p<0.005) at LacO sites, compared with LacR-GFP (53BP1: n=130; BRCA1: n=157). Averages of 3-5 experiments +/- SEM. **(D-E)** Comparison of co-localization of endogenous 53BP1 (n=247) and BRCA1 (n=223) and transfected 53BP1-Flag (n=111) and BRCA1-Flag (n=146) with LacR-SIRT6 GFP at LacO sites, assessed by IF. **(F)** Schematic representation of the Tethering assay with inhibition of signaling using Wortmannin. **(G)** co-localization of LacR-SIRT6 GFP or LacR-GFP with ɤH2AX after 24h treatment with 0µM (n[SIRT6]=43, n[GFP]=41), 1µM(n[SIRT6]=42, n[GFP]=42) or 10µM (n[SIRT6]=42, n[GFP]=41) of Wortmannin. Average of 4 experiments +/- SEM. **(H)** co-localization of LacR-SIRT6 GFP or LacR-GFP with 53BP1 upon 24h Wortmannin treatment (0µM: n[SIRT6]=32, n[GFP]=32, 1µM: n[SIRT6]=30, n[GFP]=32, 10µM: n[SIRT6]=33, n[GFP]=31). Average of 3 experiments +/- SEM. **(I)** Co-localization of LacR-SIRT6 GFP or LacR-GFP with BRCA1 upon 24h Wortmannin treatment (0µM: n[SIRT6]=30, n[GFP]=32, 1µM: n[SIRT6]=31, n[GFP]=31, 10µM: n[SIRT6]=32, n[GFP]=31). Average of 3 experiments +/- SEM. **(J)** Co-localiztion of LacR-SIRT6-Cherry or LacR-Cherry with Ku80-Flag (0µM: n[SIRT6]=45, n[Cherry]=53, 10µM: n[SIRT6]=45, n[Cherry]=58) or MRE11-Flag (0µM: n[SIRT6]=50, n[Cherry]=56, 10µM: n[SIRT6]=60, n[Cherry]=51) after 24h with or without Wortmannin. Average of 3 experiments +/- SEM. (*=p<0.05, **=p<0.005, ***=p<0.0005, ****=p<0.00005).

